# Intrinsic and non-cell autonomous roles for a neurodevelopmental syndrome-linked transcription factor

**DOI:** 10.64898/2025.12.23.696256

**Authors:** Jayson J. Smith, Seth R. Taylor, Honorine Destain, Grace Kim, David H. Hall, John G. White, David M. Miller, Paschalis Kratsios

**Affiliations:** Department of Neurobiology, University of Chicago, Chicago, IL, 60637, USA; University of Chicago Neuroscience Institute, Chicago, IL, 60637, USA; Department of Cell and Developmental Biology, Vanderbilt University School of Medicine, Nashville, TN, 37240, USA; Department of Cell Biology and Physiology, Brigham Young University, Provo, UT, 84602, USA; Department of Neuroscience, Albert Einstein College of Medicine, NY, 10461, USA; MRC Laboratory of Molecular Biology, Cambridge, UK; Program in Neuroscience, Vanderbilt University, Nashville, TN, 37240, USA

## Abstract

Transcription factors (TFs) are essential for neuronal identity, yet their potential non–cell-autonomous functions remain largely unexplored. Here, we uncover both cell- and non–cell-autonomous roles for the conserved terminal selector UNC-3 in *C. elegans* motor neurons (MNs). UNC-3 is an ortholog of human EBF3, mutations in which cause a severe neurodevelopmental syndrome. Single-cell RNA sequencing of cholinergic MNs, which express *unc-3*, and downstream GABA MNs, which do not, revealed that *unc-3* loss disrupts neuronal identity in distinct ways across MN classes. Four cholinergic MN classes lose their molecular identity entirely, whereas the AS class retains it partially, illuminating terminal selector–driven neuronal diversification processes. Integrated transcriptomic and genomic analyses uncovered a dual cell-autonomous role for UNC-3 as both a direct activator and repressor of neuron-type– specific genes in cholinergic MNs, including repression of alternate neurotransmitter programs. Unexpectedly, *unc-3* loss also caused widespread transcriptional, morphological, and connectivity defects in downstream GABA MNs. Mechanistically, these non-cell-autonomous effects are mediated by cholinergic neurotransmission and include activation of the pro-regenerative bZIP TF CEBP-1 (C/EBP) and dysregulation of UNC-6/Netrin signaling. These findings redefine terminal selectors as both intrinsic and extrinsic regulators of neuronal identity and circuit assembly, offering a mechanistic framework for understanding EBF3 syndrome pathogenesis.

## INTRODUCTION

Studies of transcription factors (TFs) have typically focused on their intrinsic (cell-autonomous) roles in guiding the development and function of cell types that produce them, herein referred to as “host” cell types. This host cell-centric framework has shaped much of our understanding of TF biology, with limited attention to whether and how TFs exert influence beyond the cells in which they are expressed. As such, the extrinsic (non-cell-autonomous) functions of TFs remain poorly understood, despite their potential to impact tissue organization, intercellular communication, and human disease. Terminal selector TFs epitomize this focus on intrinsic host cell regulation.

Defined initially in the nematode *Caenorhabditis elegans*, terminal selectors are a specialized class of TFs that orchestrates the establishment and lifelong maintenance of neuron type-specific identities^1–3^. Extensive work in *C. elegans* has revealed insights into how terminal selectors such as UNC-3/EBF1-4^4–9^, UNC-30/PITX^10–13^, and UNC-86/POU4F1-3^14,15^ activate effector genes (e.g., neurotransmitter [NT] biosynthesis proteins, NT receptors, ion channels, and neuropeptides), which determine the identity and function of their host neuron types (reviewed in^2,16–19^). To date, dozens of terminal selectors have been described for individual sensory, interneuron, and motor neuron (MN) types in the nervous system of nematodes (*C. elegans*), fruit flies (*Drosophila)*, planarians (*Schmidtea mediterranea*), cnidarians (*Nematostella vectensis)*, marine chordates (*Ciona robusta)*, zebrafish (*Danio rerio*), and mice (*Mus musculus*)^15,18,20–27^ (reviewed in^18^), representing a deeply conserved regulatory strategy for the control of neuron type identity in metazoans. Studies across model systems continue to focus on the intrinsic roles of terminal selectors in their host neuron types^28–31^. Yet, whether and how these TFs regulate the identity or properties of downstream, synaptically-connected neurons remain vastly understudied questions in the field.

UNC-3, the sole *C. elegans* ortholog of the highly conserved Collier/Olf/EBF (COE) family of TFs, functions as a terminal selector in the ventral nerve cord by controlling the identity of cholinergic MNs^4–9^, defined herein as ‘host’ cells. UNC-3 is known to act as a direct transcriptional activator in these cells^4,8^, which are further subdivided into five classes based on molecular^32,33^, anatomical and functional criteria^34,35^. However, the global effects of UNC-3 on the molecular identity of each MN class remain unknown, as only a handful of UNC-3 target genes per MN class have been identified to date using candidate approaches^4,6,7^. Notably, prior work showed that *unc-3* activity in cholinergic MNs influences synaptic connectivity in downstream GABA MNs^36,37^ but the extent to which *unc-3* non-cell-autonomously impacts the development of these cells is unknown.

Loss-of-function mutations in the *unc-3* human ortholog, *EBF3*, cause Hypotonia, Ataxia, and Delayed Development Syndrome (HADDS), also known as *EBF3* Syndrome, a recently identified neurodevelopmental disorder characterized by motor dysfunction, muscle hypotonia, intellectual disability, and autism spectrum features^38–45^. These broad and complex neurological phenotypes suggest that EBF3 plays intricate roles in both the central and peripheral nervous systems. However, our mechanistic understanding of the *EBF3* syndrome remains limited as it is largely based on clinical and *in vitro* data. Further, although cell-autonomous roles for Ebf3 have been described in cortical neuron migration in mice^46,47^, it’s non-cell-autonomous roles remain unknown. Like *unc-3* in *C. elegans*, *EBF3* is expressed in human MNs^48–50^. In addition, UNC-3 exhibits high amino-acid sequence similarity and conserved domain structure with EBF3 (**Fig. 1A**). Therefore, a detailed analysis of UNC-3’s regulatory roles in *C. elegans*, encompassing effects both in host cells and downstream MNs, promises to not only advance our understanding of TF biology, but could also generate early insight into the *EBF3* neurodevelopmental syndrome.

**Figure 1.**
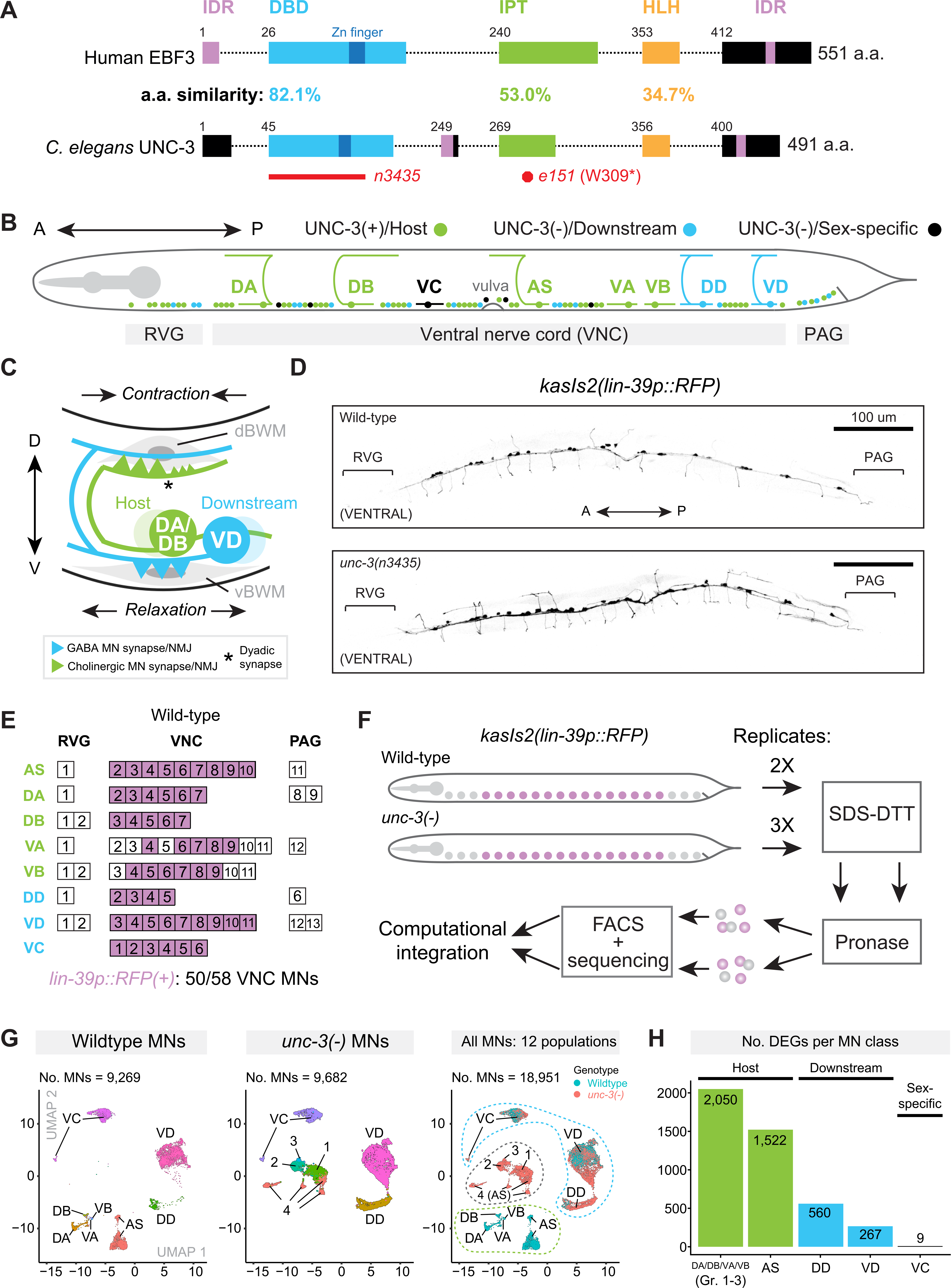
Single-cell profiling of host and downstream MNs in the presence and absence of UNC-3/EBF. **(A)** Schematic of human EBF3 and *C. elegans* UNC-3 proteins. Red bar below UNC-3 shows region deleted in the *unc-3(n3435)* allele used for scRNA-seq and all genetic experiments in this study. Red octagon indicates the location of the *unc-3(e151)* nonsense mutation used for connectivity analyses in Figure 8C-E. Conserved protein domains are listed above UNC-3 and amino acid (a.a.) similarity is indicated as a percentage between protein schematics. DBD, DNA binding domain (light blue); IDR, Intrinsically Disordered Region (pink); IPT, Immunoglobulin-like/Plexin/Transcription Factor (green); HLH, Helix-Loop-Helix (orange). (**B)** Schematic of host (UNC-3[+], green), downstream (UNC-3[-], blue), and sex-specific (UNC-3[-], black) MNs in the *C. elegans* VNC. RVG, Retrovesicular Ganglion; PAG, Preanal Ganglion. **(C)** Segment of the *C. elegans* motor circuit. d/vBWM, dorsal/ventral Body Wall Muscle. NMJ, Neuromuscular Junction. **(D)** *lin-39p::RFP* reporter expression in WT (top) and *unc-3(-)* mutants (bottom). **(E)** Single-cell expression pattern of *lin-39p::RFP* reporter in WT Day 1 young adult VNC MNs. **(F)** ScRNA-seq strategy used to profile adult VNC MNs. **(G)** UMAPs showing sequenced MNs derived from WT only (left), *unc-3(-)* mutant only (middle), or all MNs (right). Left and middle panels are colored by MN class; right panel is colored by genotype of origin. Numbers (i.e., 1-4) refer to mutant MN groups lacking clear wild-type counterparts. In right panel, green dotted line indicates the five wild-type-specific MN populations, blue dotted line indicates the three MN populations which are derived from both genotypes, and the grey dotted line indicates the four *unc-3(-)*-specific MN populations. **(H)** Bar plot depicting the total number of DEGs identified in each MN class.

Here, we leverage the powerful genetics and fully mapped nervous system^34,51–54^ of *C. elegans* to uncover the effects of *unc-3* loss not only in cholinergic (host) MNs, but also in GABA MNs, which do not express *unc-3*^32,33^, but receive cholinergic input from host MNs (**Fig. 1B-C**). To this end, we employ single-cell RNA sequencing (scRNA-seq) to identify global transcriptional changes in both MN populations. In the five classes of host MNs, loss of the same TF (*unc-3*) leads to non-uniform effects: One class partially retains its original identity, whereas the remaining four collapse into three mutant cell populations, an important finding for our understanding of neuronal subtype diversification (see Discussion) that would have been masked in a bulk RNA-seq approach. In host MNs, we uncovered a dual role for UNC-3 as both a transcriptional activator and major repressor of gene expression. In downstream GABA MNs, we show that UNC-3 exerts extensive non-cell autonomous effects by altering their transcriptome, morphology, and connectivity. Mechanistically, these effects are mediated by *unc-3*-dependent cholinergic neurotransmission from host to GABA MNs and include activation of the conserved neuronal regeneration-related TF CEBP-1 (C/EBP), as well as Netrin signaling dysregulation. Our study reveals critical roles for UNC-3/EBF in shaping both host neuron identity and downstream neuron development, offering a conceptual framework for understanding cell- and non-cell autonomous functions of TFs that could inform human disease pathogenesis.

## RESULTS

### The *C. elegans* ventral nerve cord as a model for studies of TF functions in host and downstream neurons

The adult *C. elegans* hermaphrodite contains 302 neurons^51^. UNC-3 functions as a terminal selector in 50 cholinergic MNs (‘host’ cells) located in the ventral nerve cord (VNC), a structure analogous to the vertebrate spinal cord^51^, and its flanking retrovesicular and preanal ganglia (RVG, PAG; **Fig. 1B**). An additional 19 GABA and six egg-laying MNs are in the VNC, RVG, and PAG, and do not express *unc-3*^32,33^. All these MNs are categorized into eight distinct classes based on morphological and functional criteria (**Fig. 1B**). Of the five classes of cholinergic host MNs: two control forward locomotion (DB, VB)^35^, two control backward locomotion (DA, VA)^35^, and one class (AS) contributes to anterior-posterior bending wave propagation^55^. In addition, two classes (DD, VD) of GABA MNs receive cholinergic input by host MNs^54^, and are referred to here as ‘downstream MNs’. Last, the VC class is cholinergic and controls egg-laying^34^(**Fig. 1B**). These eight classes subdivide further into at least 29 molecularly distinct subclasses based on single-cell transcriptomic profiling^32,33^. The coordinated activity of cholinergic host MNs and GABA downstream MNs propels the characteristic sinusoidal locomotion (i.e., undulation) of *C. elegan*s^34,35,51^. Cholinergic MNs are connected with GABA MNs and muscle via dyadic synapses (**Fig. 1C**) ^35^. To generate dorsal body bends, DA/DB classes of cholinergic MNs simultaneously stimulate dorsal muscle to contract and downstream GABA MNs (VD class), which in turn relax ventral muscle^35^ (**Fig. 1C**). Conversely, to generate ventral body bends, VA/VB cholinergic MNs stimulate ventral muscle and a different class of GABA MNs (DD class)^35^. Last, the lineage and connectivity of all these MNs have been mapped^34,51^, offering a well-characterized *in vivo* system to dissect the roles of UNC-3 both in host and downstream cells at high molecular and cellular resolution.

### Loss of *unc-3* differentially affects the transcriptome of both host and downstream cells

To identify the functions of UNC-3 in both cholinergic (host) and GABA (downstream) cells, we performed scRNA-seq on MNs isolated from day 1 adult hermaphrodites of wild-type (WT) and *unc-3(n3435)* null mutant (*unc-3(-)*, **Fig. 1A**) animals with two and three biological replicates, respectively. MNs were labeled with a *lin-39p::RFP* reporter, which marks 86% (50/58) of host and downstream MNs of the VNC and is unaffected by the loss of *unc-3* (**Fig. 1D-E**). Cells were isolated through FACS and sequenced following established protocols^32,33,56^ (**Fig. 1F**). Upon quality control (see Methods), 9,269 WT and 9,682 *unc-3(-)* MN transcriptomes were obtained across biological replicates (**File S1)**.

Given the striking molecular diversity that we recently uncovered among adult VNC MNs of WT animals^32,33^, at least two hypothetical outcomes can be envisioned for the five cholinergic host MN classes (DA, DB, VA, VB, AS) lacking *unc-3*: a uniform collapse of identity into a single undifferentiated state (i.e., erasure of class identities) or MN class-specific effects with varied degrees of identity loss. Similarly, for the VNC MNs that do not express *unc-3* - downstream MN classes (DD, VD) and the VC class - at least three outcomes can be predicted in the *unc-3(-)* mutant: no effect, uniform transcriptional disruption, or class-specific effects.

To assess these possibilities, we performed computational integration and UMAP clustering of cells from both WT and *unc-3(-)* genotypes. This approach yielded 12 transcriptionally distinct MN populations, consisting of five WT-specific populations, four *unc-3(-)-*specific populations, and three populations shared between the two genotypes (**Fig. 1G**). By integrating differential expression analysis with known MN class-specific markers (**File S1, Fig. S1A**), we found that the five WT-specific clusters precisely corresponded to the five host MN classes (AS, DA, DB, VA, VB), whereas the three genotype-shared clusters express molecular markers of the two downstream GABA classes (DD, VD) and the VC class (**Fig. 1G, File S1)**. Notably, the four *unc-3(-)*-specific MN populations (Groups 1–4) lacked direct WT MN class or subclass counterparts, with only Group 4 exhibiting partial transcriptomic similarity to wild-type AS MNs (**Fig. 1G, File S1**). As the *lin-39p::RFP* reporter remains mostly restricted to VNC MNs in the *unc-3(-)* background (**Fig. 1D-E**), we reasoned that these four mutant populations represent host MN classes deprived of their terminal selector TF UNC-3.

Next, we assessed the number of differentially expressed genes (DEGs) in each class as a metric for transcriptional disruption. Because Groups 1-3 lack class-specific gene expression, we combined the transcriptomes of DA, DB, VA, and VB WT MNs to compare collectively to Groups 1-3. We found that these 4 classes of host cholinergic MNs had the most DEGs (2,050), exhibiting apparent collapse in their molecular identity, followed by the 5^th^ host class - AS MNs - which had 1,522 DEGs (discussed below, **Fig. 1H**). Surprisingly, even though the GABA (DD, VD) and VC MN classes do not express *unc-3*^32,33^ and overlap in UMAP space between genotypes (**Fig. 1G, S1B-C**), we found that the DD and VD neurons underwent significant transcriptomic disruption in the *unc-3(-)* mutant with 560 and 267 DEGs between genotypes, respectively, whereas the VC MNs were essentially unchanged (9 DEGs) (**Fig. 1H**). We conclude that loss of *unc-3* differentially affects the transcriptomes of host and downstream MNs.

### AS MNs are partially resilient to *unc-3* loss

The single-cell resolution of our molecular profiling approach revealed non-uniform effects of *unc-3* loss on the five host MN classes: DA, DB, VA, and VB MNs lose their original identity, whereas AS MNs partially retain their molecular identity (**Fig. 2A**). To probe this further, we used NeuroPAL^57^, a multicolor strain suitable for nervous system-wide cell identification and detection of neuronal identity changes based on fluorescent marker expression. Indeed, we found that AS MNs (green) as well as downstream (DD, VD) (blue) and VC (unlabeled) MNs retained their fluorescent color codes in *unc-3(-)* animals, whereas DA, DB, VA, and VB host MNs instead adopt a uniform dim-red color (**Fig. 2B-C**). These results are consistent with the notion that AS MNs are uniquely resilient to loss of UNC-3 among the host cholinergic MN classes.

**Figure 2.**
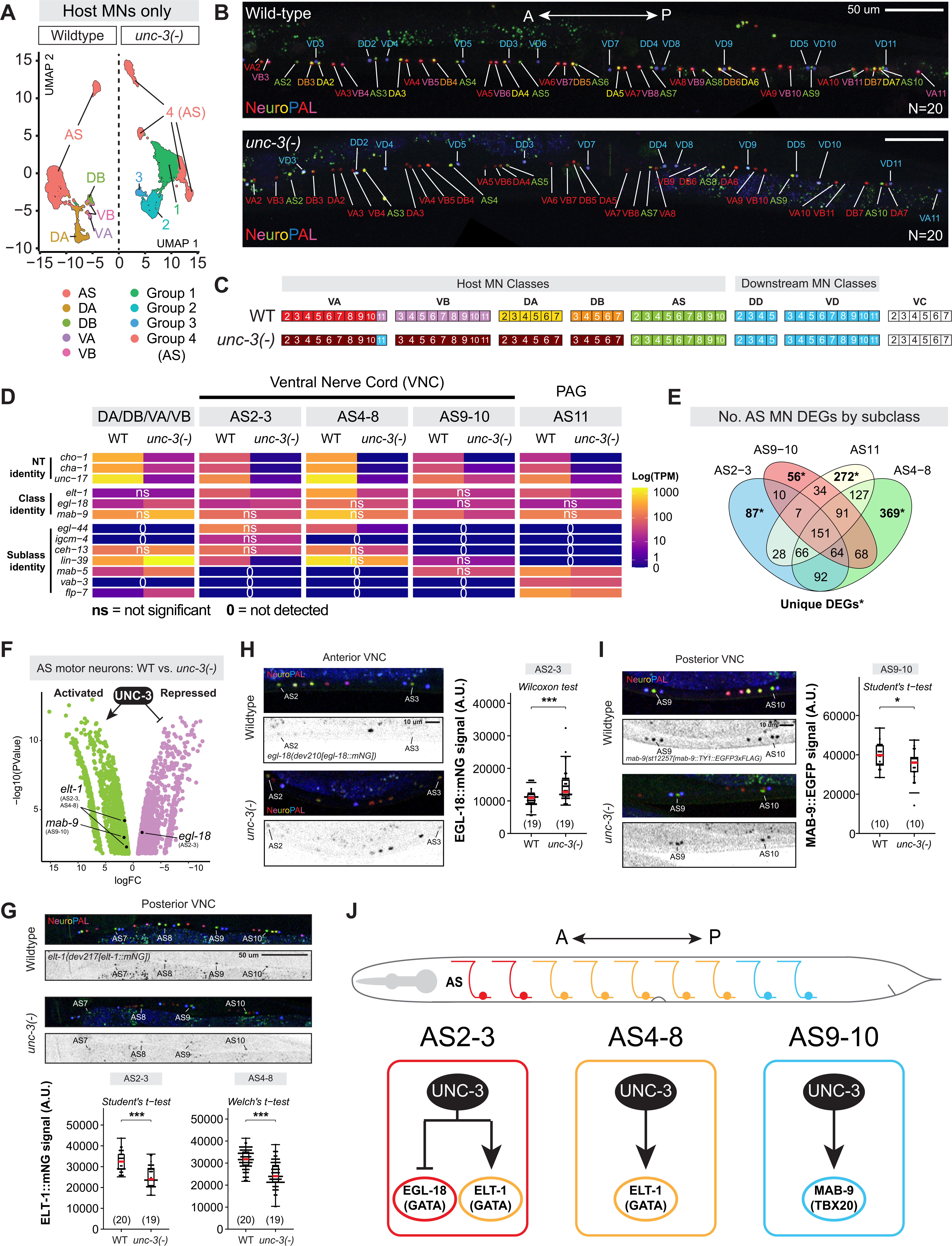
AS MN identity partially persists in the absence of UNC-3/EBF. **(A)** UMAP of integration of host MN classes in WT and *unc-3(-)* backgrounds. Clusters are colored by population identity, as indicated below the UMAP. **(B)** Representative micrographs of NeuroPAL transgene in VNC MNs in WT (top) or *unc-3(-)* (bottom) animals. **(C)** Table representation of NeuroPAL-based color codes from B. **(D)** Heatmap of log expression (Transcripts per million, TPM) of cholinergic identity genes, AS class identity genes, and AS subclass identity genes in WT vs. *unc-3(-)* host MNs. ns, not significant; 0, not detected. **(E)** Venn diagram depicting number of shared and unique DEGs across AS MN subclasses. **(F)** Volcano plot of all significant DEGs (False Discovery Rate, FDR < 0.05) across AS MN subtypes in *unc-3(-)* MNs relative to WT. Green, UNC-3-activated; Pink, UNC-3-repressed. **(G-I)** Fluorescence micrographs of NeuroPAL (top) and endogenous fluorescent reporters (bottom) for TFs ELT-1 (G), EGL-18 (H), and MAB-9 (I) with quantifications of fluorescence intensity (A.U., arbitrary units) in wild-type or *unc-3(-)* day 1 adult animals. AS subclasses quantified are listed above each plot. Number of animals (N) is listed in each plot. Statistical tests were selected based on data distribution as follows: Normally distributed data (Shapiro-Wilk *p* > 0.05 in both groups) with equal variances (Levene’s test *p* > 0.05) = Student’s t-test; Normally distributed with unequal variances = Welch’s t-test; Non-normal data = Wilcoxon rank-sum test. For all quantifications/statistical tests, p>0.05 = not significant, ns; p<.05 = *; p<.01 = **; p<.001 = ***. Box-plot elements: thick horizontal line (red) = median; box = 25th to 75th percentiles (interquartile range); whiskers extend to the furthest data point within 1.5 × IQR from the quartiles (or the min/max if all points lie within this range); individual data points are overlaid as black dots. **(J)** Summary of *in vivo* validated AS MN subclass-specific effects of UNC-3 on TFs quantified in G-I.

In WT animals, the AS class is subdivided into four molecularly distinct subclasses: AS2–3, AS4–8, AS9–10, and AS11^33^. Here, we observed that loss of *unc-3* leads to the formation of four distinct mutant AS subclusters (**Fig. 2A**), suggesting that each of the four WT subclass identities might be partially retained. To test this idea, we assessed the expression of Hox genes, which delineate each AS subclass along the A-P axis of the VNC^33^. One mutant AS subcluster had an anterior Hox profile (*ceh-13/HOXB1/HOXD1[+]*, *mab-5/HOXB8/HOXC8[-]*) indicating AS2-3 subclass identity, supported by its co-expression of *egl-44/TEAD* and *igcm-4*/*FGFRL1*, which uniquely label this subclass in WT^33^ (**Fig. 2D**). Similarly, the anterior-midbody Hox gene *lin-39/HOXA5* was highly enriched in one subcluster lacking *mab-5*, matching the AS4-8 subclass^33^ (**Fig. 2D**). Only one subcluster expressed both *lin-39* and *mab-5*, consistent with the WT AS9-10 posterior-midbody subclass^33^ (**Fig. 2D**). The last AS subcluster had the expected posterior Hox gene expression profile of AS11 (*lin-39[-], mab-5[+]*), supported by co-expression of *vab-3/PAX6* and *flp-7*^33^ (**Fig. 2D**). Altogether, this analysis mapped the *unc-3(-)* AS MN subclasses to their WT counterparts, suggesting that AS neuron identity partially persists in the absence of *unc-3*.

Supporting this idea, DEG analysis followed by reporter validation revealed partial effects of *unc-3* loss in each AS subclass (**Fig. 2E-F**). For example, the GATA1/3 TF *elt-1* and the vesicular acetylcholine (ACh) transporter *unc-17/VAChT* are both normally expressed in all AS neurons. In *unc-3(-)* mutants, they show partial reductions in anterior AS2-3 (*elt-1*: |log2FC| = 1.2; *unc-17*: |log2FC| = 5.4) and midbody AS4-8 subclasses (*elt-1*: |log2FC| = 1.6; *unc-17*: |log2FC| = 6.8), but remain unaffected (*elt-1*) or only moderately reduced (*unc-17*) in the posterior AS9-10 subclass (**Fig. 2D, F**). We independently confirmed these effects in *unc-3(-)* mutants using an endogenous *elt-1* reporter (**Fig. 2G**). Similarly, the anterior-midbody HOX TF *lin-39*, which is expressed in both anterior and midbody AS subclasses of WT animals, is significantly reduced (|log2FC| = 5.8) only in the anterior AS2-3 subclass of *unc-3(-)* animals. To further corroborate these findings, we used endogenous fluorescent reporters and NeuroPAL to validate UNC-3 repression of the TF *egl-18/GATA4* in AS2-3 (**Fig. 2H**) and its activation of *mab-9/TBX20* in AS9-10 (**Fig. 2I**). Altogether, scRNA-seq and endogenous reporter data support a model where UNC-3 contributes to AS subclass identity by regulating distinct TFs (**Fig. 2J**).

### UNC-3/EBF acts both as a transcriptional activator and repressor in host MNs

Prior work on UNC-3 identified a limited number of target genes in cholinergic host MNs following candidate gene approaches, supporting its role as a direct transcriptional activator^4,6–8^. To uncover the global effects of *unc-3* loss in host MNs in an unbiased manner, we performed pseudobulk comparisons between WT and *unc-3(-)* MNs. Because the transcriptional effects on AS MNs were described in detail above, we focused here on the remaining four classes (DA, DB, VA, VB) of host MNs (**Fig. 2A**). In total, we identified 929 downregulated transcripts in *unc-3(-)* MNs (False Discovery Rate [FDR] < 0.05), including 49 target genes known to be activated by UNC-3, such as ACh biosynthesis components (e.g., *cha-1/ChAT*, *cho-1/ChT*, *unc-17/VAChT*)^4^, MN class-specific TFs (e.g., *bnc-1/BNC1-2*^7,8^, *ceh-12/MNX1*^4^, *unc-4/UNCX*^4,58^, *vab-7/EVX1-2*^4^), ion channels (e.g., *nca-1/NALCN*^4^*, lgc-36/GABRA1-3*^7^*, slo-2/KCNT1-2*^9^), and neuropeptides (e.g., *nlp-21*^4^*, flp-18*^6^) (**Fig. 3A**, **S2A**). Strikingly, we identified even more upregulated transcripts (1,121) in *unc-3(-)* host MNs, including six known UNC-3-repressed target genes (*glr-5/GRIK4*^5,9^, *lin-11/LHX1*^58^*, ida-1/PTPRN2*^9,58^*, ser-2/HTR1A*^9^*, lin-39/HOXA5*^59^*, mab-5/HOX8*^59^). These results, supported by the validation of two novel repressed targets (*egl-18/GATA4*, **Fig. 2H**; *ceh-6/POU3F*, **Fig. S2E**) using endogenous fluorescent reporters, suggest a major role for UNC-3 in gene repression.

**Figure 3.**
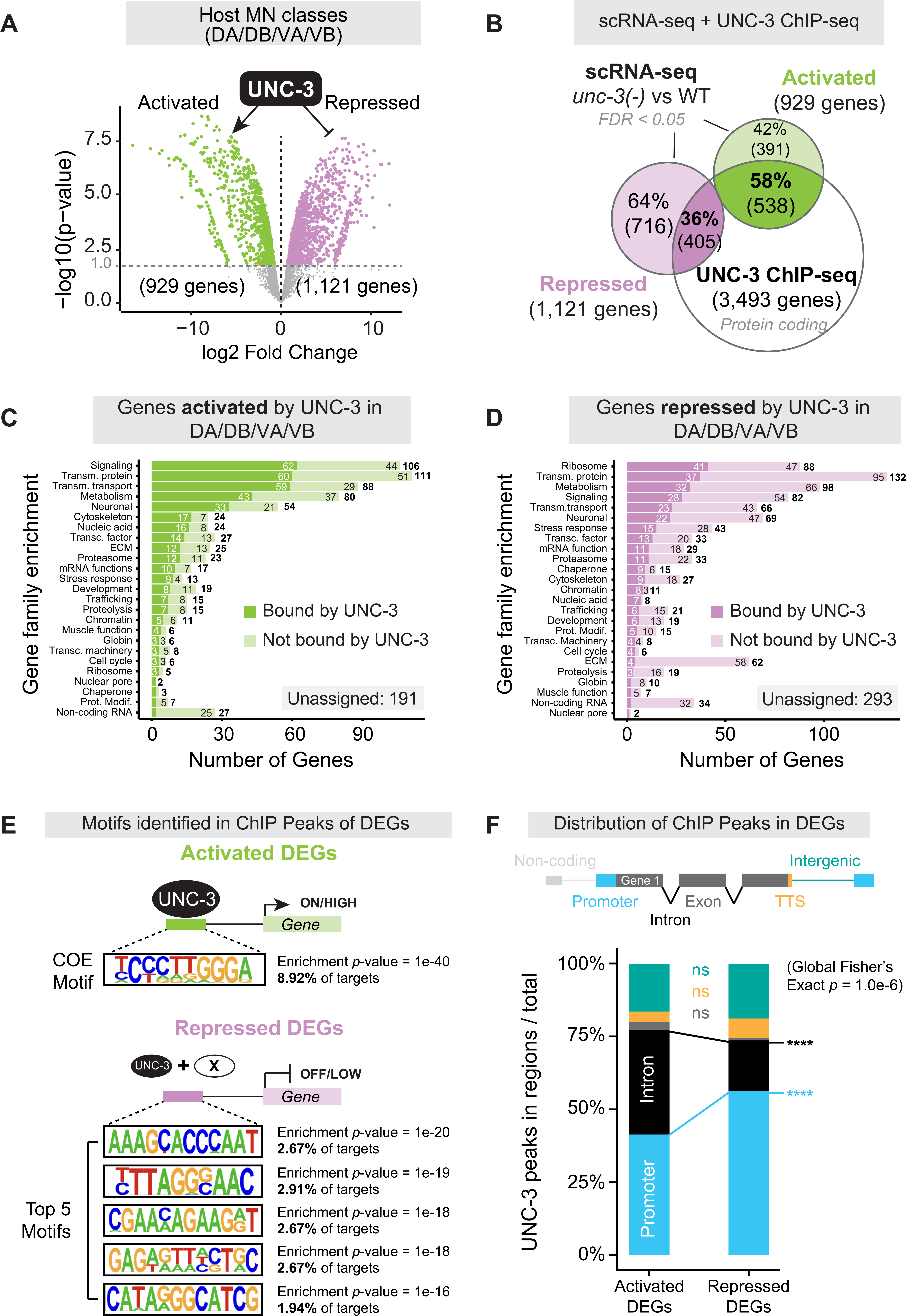
UNC-3/EBF is a dual transcriptional regulator in host MNs. **(A)** Volcano plot of results from pseudobulk analysis showing all significant DEGs (False Discovery Rate, FDR < 0.05) in DA/DB/VA/VB MNs of WT vs. *unc-3(-)*. Green dot = UNC-3-activated gene; Pink dot = UNC-3-repressed genes. **(B)** Venn diagram depicting the integration of DEGs from scRNA-seq with UNC-3 ChIP-seq data^8^. **(C-D)** Bar charts depicting WormCat2^60^ gene family enrichment of activated (C) or repressed (D) DEGs in host MNs overlayed with UNC-3 ChIP-seq data, as in (B). **(E)** Schematic of most significantly enriched motifs within UNC-3 ChIP-seq peaks in cis-regulatory loci of activated (top) or repressed (bottom) DEGs. The most significant motif for activated DEGs was UNC-3’s cognate COE binding motif (shown), whereas the top five motifs present in repressed DEGs are shown. Enrichment *p*-value and proportion of targets with indicated motif are listed to the right of each sequence. **(F)** Top: schematic of a generic gene locus with relevant regions compared below. Bottom: proportions of UNC-3 ChIP-seq peaks in each genomic region were compared between activated and repressed DEGs using Fisher’s exact test (global) followed by post-hoc 2×2 Fisher’s exact tests with Bonferroni correction. p>.05 = Not significant, ns; p<.05 = *; p<.01 = **; p<.001 = ***; p<.0001 = ****. Colors of asterisk/ns indicate respective categories compared in post-hoc tests.

Gene enrichment analysis (WormCat2.0)^60^ of activated and repressed targets revealed categories related to neuronal function, such as synaptic signaling, membrane transport, and metabolism (**Fig. 3C-D, S2B-D, Files S3-4**). To determine whether the widespread effects of *unc-3* loss on the transcriptomes of host MNs are direct or indirect, we integrated our scRNA-seq data with available UNC-3 ChIP-seq data^8^. We found UNC-3 ChIP-seq peaks at the *cis*-regulatory region of 58% of activated and 36% of repressed target genes (**Fig. 3B**), indicating that UNC-3 can act directly to either activate or repress gene expression in host MNs. Altogether, the unbiased identification of UNC-3 target genes in host MNs supports a dual role. Based on the 929 downregulated genes most of which are UNC-3-bound (58%), UNC-3 is a critical determinant of MN identity by acting chiefly as a direct transcriptional activator, which is consistent with prior work^4,6–8^. Further, the identification of 1,121 upregulated genes points to a new major role for UNC-3 as transcriptional repressor in host MNs. Interestingly, only 36% of these are UNC-3-bound, suggesting that UNC-3 primarily acts indirectly to repress gene expression.

### UNC-3/EBF represses targets via non-cognate motifs and promoter-biased binding

Prior work in host MNs showed that UNC-3 directly activates dozens of terminal identity genes via its cognate DNA binding site (TCCCNNGGA)^4,5,8^, termed COE motif. However, the precise mechanisms through which UNC-3 represses gene expression in host MNs remain poorly understood. The integration of scRNA-Seq and ChIP-Seq data suggests that UNC-3 can act indirectly or directly to repress gene expression (**Fig. 3B**). To investigate the latter at the genomic level, we performed motif enrichment analysis (HOMER, see Methods) on sequences contained within UNC-3 ChIP-seq peaks in activated or repressed DEGs in host MNs. The COE motif was the most enriched sequence within UNC-3 ChIP-seq peaks of activated target genes (**Fig. 3E, File S5**), corroborating a direct role for UNC-3 in transcriptional activation^4,5,8^. Surprisingly, however, our analysis did not recover the COE motif in UNC-3 peaks of repressed targets (**Fig. 3E, File S6**), indicating UNC-3 primarily binds non-cognate motifs to directly repress its target genes. We independently validated this finding using TargetOrtho2^61–63^, which identified COE motifs in 61.3% (330/538 genes) of UNC-3-bound activated targets but only 24.4% (99/405) of UNC-3-bound repressed targets (**Fig. S4, File S7**). To gain further insight into its repressive mechanism, we examined the genomic distribution of UNC-3 ChIP-seq peaks in the *cis*-regulatory regions of DEGs. UNC-3 equally occupied introns and promoters in activated targets but showed a strong preference for promoters in repressed targets (**Fig. 3F**). Thus, we propose that the direct repressor role of UNC-3 differs from its activator function by largely excluding the cognate COE motif and exhibiting a promoter bias.

### UNC-3/EBF represses alternate neurotransmitter identity genes in host MNs

To further probe the repressor role of UNC-3 in host MNs, we focused on gene enrichment categories of UNC-3-repressed targets (**Fig. 3D**). We found an overrepresentation of neuronal and cell-cell signaling-related categories, including neuropeptides, G protein-coupled receptors, and homeodomain TFs (**Fig. S2B, D**). Strikingly, we also observed NT identity genes normally restricted to other neuron classes. These included GABA (*unc-46/LAMP*, *unc-47/VGAT*), glutamatergic (*eat-4/VGLUT*), and monoaminergic (*cat-1/VMAT*, *tph-1/TPH1-2*) identity genes (**Fig. 4A**). We independently validated the ectopic expression of these five genes in *unc-3(-)* host MNs using fluorescent reporters (**Fig. 4B, S5A-E**).

**Figure 4.**
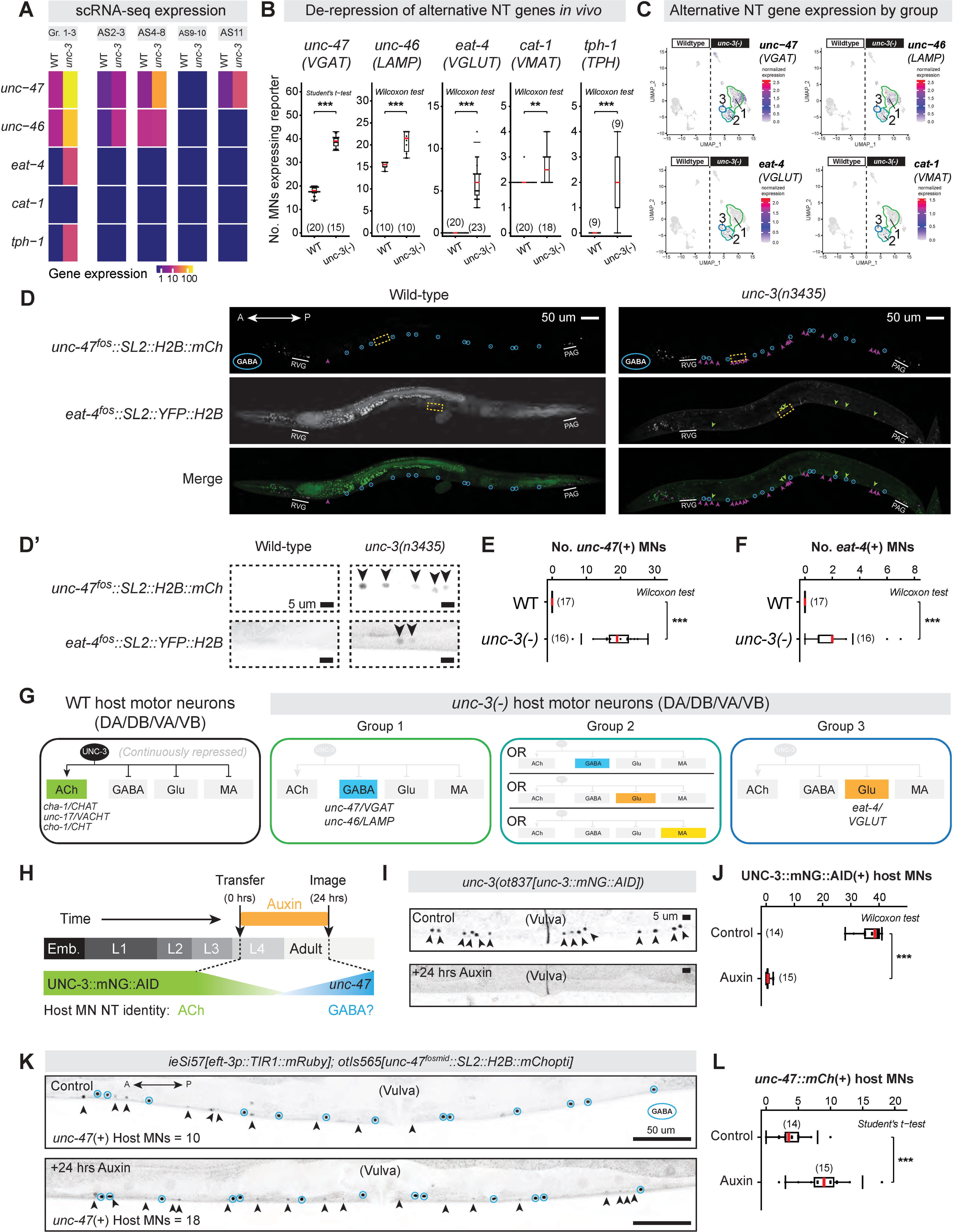
UNC-3/EBF represses alternative neuronal identities in host MNs. **(A)** Heatmap depicting differential expression (Transcripts per million, TPM) of alternative neurotransmitter (NT) identity genes in WT vs. *unc-3(-)* MNs. **(B)** Quantification of host MNs expressing fluorescent reporters for alternative NT identity genes *in vivo*. **(C)** Feature plots showing that ectopic expression of alternative NT identity genes is non-uniform across *unc-3(-)* mutant MN groups 1-3. **(D)** Fluorescence micrographs of *unc-47* (GABA) and *eat-4* (glutamatergic) reporters in WT or *unc-3(-)* animals. Magenta and green arrowheads indicate host MNs ectopically expressing *unc-47* or *eat-4*, respectively. Downstream/GABA MNs (DDs/VDs) are circled in blue. **(D’)** Insets of *unc-47* (top) or *eat-4* (bottom) fluorescence expression from yellow boxes in D. **(E-F)** Quantification of host MNs ectopically expressing the *unc-47* (E) or *eat-4* (F) fluorescent reporter. **(G)** Schematic summary of UNC-3’s role as a dual transcriptional regulator in host MNs. **(H)** Strategy for temporal UNC-3 depletion coupled with quantification of ectopic *unc-47* expression in host MNs. **(I)** Representative micrographs of UNC-3::mNG::AID expression in control (top) or 24-hour auxin-treated (bottom) animals. **(J)** Quantification of host MNs expressing UNC-3::mNG::AID in control or 24-hour auxin-treated animals. **(K)** Fluorescence micrographs of *unc-47* in control (top) or 24-hour auxin-treated (bottom) animals. **(L)** Quantification of host MNs ectopically expressing *unc-47* in control or 24-hour auxin-treated animals. All quantifications were performed at young adult stage. Number of animals (N) is listed in each plot. Statistical tests were selected based on data distribution as follows: Normally distributed data (Shapiro-Wilk *p* > 0.05 in both groups) with equal variances (Levene’s test *p* > 0.05) = Student’s t-test; normally distributed with unequal variances = Welch’s t-test; non-normal data or zero expression = Wilcoxon rank-sum test. For all quantifications/statistical tests, p>.05 = Not significant, ns; p<.05 = *; p<.01 = **; p<.001 = ***. Box-plot elements: thick horizontal line (red) = median; box = 25th to 75th percentiles (interquartile range); whiskers extend to the furthest data point within 1.5 × IQR from the quartiles (or the min/max if all points lie within this range); individual data points are overlaid as black dots.

Next, we asked whether host MNs in *unc-3(-)* animals ectopically co-express multiple alternate NT identity genes (e.g., dual NT identity), or if they are subdivided into groups each expressing a distinct NT identity. To this end, we examined individual *unc-3(-)* MNs for co-expression of GABA (*unc-46, unc-47*), glutamatergic (*eat-4*), and monoaminergic (*cat-1*) identity markers. We noted that Group 1 expresses *unc-47* and *unc-46,* but lacks *eat-4* and *cat-1* (**Fig. 4C**). Consistently, animals that simultaneously carry a red (mChOpti) reporter for *unc-47/VGAT* and a yellow (YFP) reporter for *eat-4/VGLUT* did not show any overlapping expression in individual *unc-3(-)* host MNs (**Fig. 4D-F, S5F-G**). Using a similar analysis, we identified two additional groups. Group 2 contains cells that either express glutamatergic (*eat-4*), GABA (*unc-46, unc-47*), or monoaminergic (*cat-1*) markers, whereas Group 3 only expresses low levels of the *eat-4* glutamatergic marker. Altogether, the single-cell resolution of RNA-seq revealed that, in the absence of UNC-3, host MNs are divided into several groups, each adopting molecular features indicative of a distinct NT identity: MNs of Group 1 acquire GABA features, MNs of Group 2 can adopt features of three alternate NT identities, and MNs of Group 3 can acquire glutamatergic features (**Fig. 4G**).

To determine whether *unc-3(-)* host MNs fully or partially adopt alternate NT identities, we focused on Group 1. Although *unc-46/LAMP* and *unc-47/VGAT* are ectopically co-expressed in this group (**Fig. 4A-C**), other core GABA NT identity features, such as the biosynthetic enzyme gene *unc-25*/*GAD*, were unaffected (**Fig. S1A**). Further, *unc-30/PITX*, the terminal selector of GABA MN identity and direct activator of *unc-46* and *unc-47*^10–13^, was neither ectopically expressed in host MNs of *unc-3* mutants nor bound by UNC-3 at its *cis*-regulatory sequences (**Fig. S3B-C**). Instead, analysis of ChIP-seq^8^ and TargetOrtho2 data revealed UNC-3 binding to a non-cognate motif in the *unc-47/VGAT* promoter (**Fig. S5A**), suggesting direct transcriptional repression of this GABA NT identity gene by UNC-3. Collectively, these results argue against cell fate transformation and instead support a model in which UNC-3 represses latent alternate NT identity genes to secure the cholinergic NT identity of host MNs.

### UNC-3/EBF is continuously required to repress GABA NT identity in host MNs

UNC-3 is expressed throughout life in adult host MNs to maintain the expression of ACh biosynthesis genes^8^. To test whether UNC-3 also continuously represses alternative NT identity genes, we used the auxin-inducible degron (AID) system^64^ to acutely deplete UNC-3 in late larval/early adult stages. We started auxin treatment at larval stage 4 (L4), by which time all nerve cord MNs have differentiated, and discontinued auxin treatment at day 1 of adulthood (**Fig. 4H**). After confirming efficient UNC-3 depletion in day 1 adult MNs (**Fig. 4I-J**), we observed a mild hypomorphic effect on *unc-47/VGAT* expression in control animals (**Fig. 4E, 4L**) but robust ectopic expression in host MNs of animals treated with auxin (**Fig. 4K-L**), indicating that UNC-3 activity is required in late larval/early adult life to repress alternate NT identity genes.

### Widespread transcriptional dysregulation of downstream GABA MNs in *unc-3* mutants

In post-mitotic neurons, TFs have been typically studied within their host neuron types. Whether and how TFs control the development of downstream post-mitotic neurons remains largely unknown. To address this question, we systematically characterized the transcriptomes of GABA MNs (DD/VD) in *unc-3(-)* animals. Our scRNA-seq data show that GABA NT identity markers, such as *unc-25/GAD*, *unc-47/VGAT,* and *gta-1/GABA-T,* remain unchanged in GABA MNs of *unc-3(-)* mutants (**Fig. S6A**). Additionally, unlike host MNs, we observed no ectopic expression of alternative NT identity genes (e.g., cholinergic, glutamatergic) in GABA MNs (**File S2**). However, we observed other extensive transcriptomic changes in GABA MNs of *unc-3(-)* animals with 560 DEGs in DD MNs and 267 DEGs in VD MNs, most of which were upregulated (**Fig. 5A**). These effects appear specific to downstream GABA MNs, as we only identified 9 DEGs in VC MNs (**Fig. 5A**), which – unlike GABA MNs – do not receive any synaptic input from host MNs^34,54^. Most DEGs were specific to anterior-midbody GABA MN subclasses, with 68% (256/377 genes) of DD-specific DEGs unique to DD2-3 subclass and 92% (77/84 genes) of VD-specific DEGs unique to VD3-7 subclass (**Fig. 5B, S6B-C**). Given that UNC-3 is not expressed in the embryonic lineage of (or mature) DD or VD MNs (**Fig. S7A-B’’**)^65–67^, these results suggest a non-cell autonomous impact of *unc-3* on the transcriptomes of downstream GABA cells.

**Figure 5.**
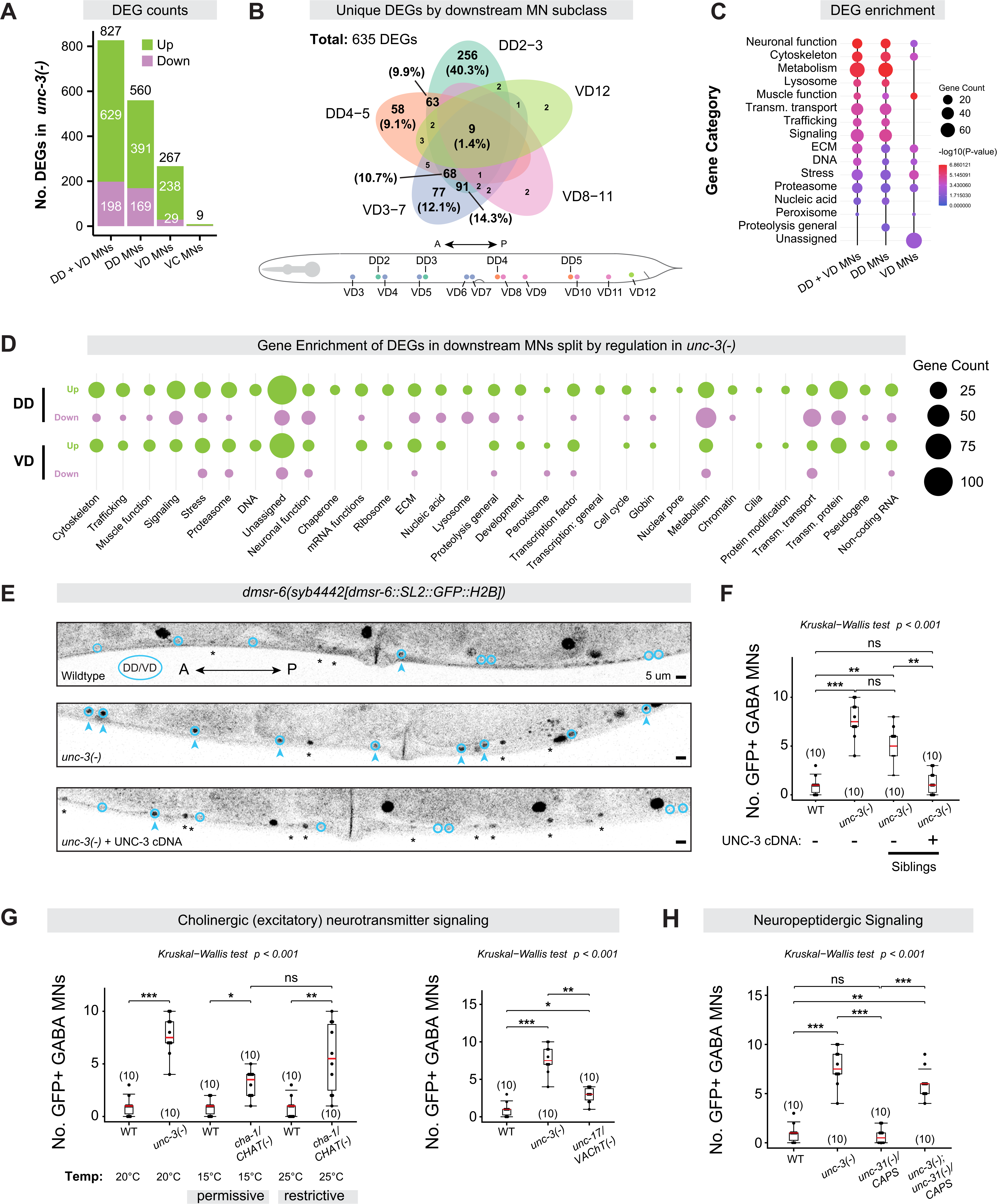
UNC-3/EBF loss leads to widespread transcriptomic disruption in downstream MNs. **(A)** Stacked bar chart depicting number of differentially expressed genes (DEGs) in non-host MN classes (DD, VD, VC). **(B)** Top: Venn diagram of shared and unique DEGs by GABA MN subclass. Numbers represent counts of unique DEGs in each subclass and percentages provide fraction of total DEGs. Bottom: *C. elegans* schematic of GABA MN subclasses color matched to Venn diagram. **(C)** Gene enrichment (WormCat 2.0^60^) analysis for DEGs in downstream (DD, VD) MNs. Dot sizes correspond to gene counts; dot colors correspond to -log_10_(p-Value). **(D)** DEG enrichment across biological categories split by regulation (upregulated = green; downregulated = pink) and MN class. Dot sizes correspond to gene counts. **(E)** Representative micrographs depicting endogenous *dmsr-6* expression in the VNC of WT, *unc-3(-)* mutant, and *unc-3(-)* mutant with UNC-3 cDNA rescue construct. Downstream/GABA MNs are circled in blue. Black asterisks and blue arrowheads indicate host (cholinergic) and downstream (GABA) MNs expressing *dmsr-6*, respectively. **(F)** Number of GABA MNs expressing *dmsr-6* in each in WT, *unc-3(-)* mutants, and *unc-3(-)* mutant siblings without and with UNC-3 cDNA rescue construct. Groups were statistically compared globally via a Kruskal-Wallis test, with *p*-values provided for significant and relevant comparisons. ns, not significant. **(G-H)** Number of VNC MNs expressing endogenous *dmsr-6* reporter in wild-type animals or animals harboring mutations in components of cholinergic synaptic neurotransmission (G) or neuropeptidergic signaling (H). All quantifications were performed at young adult stage. Number of animals (N) is listed in each plot. Kruskal-Wallis tests were performed, followed by Dunn’s post-hoc test with Benjamini-Hochberg adjustment due to non-normally distributed data (Shapiro-Wilk test in groups with n ≥ 5 and SD > 0). Global Kruskal-Wallis p-value is displayed at the top. For pairwise comparisons, p<.05 = *; p<.01 = **; p<.001 = ***. Only relevant comparisons are shown. Box-plot elements: thick horizontal line (red) = median; box = 25th to 75th percentiles (interquartile range); whiskers extend to the furthest data point within 1.5 × IQR from the quartiles (or the min/max if all points lie within this range); individual data points are overlaid as black dots.

To determine which biological processes are dysregulated in downstream (DD, VD) MNs in the absence of *unc-3*, we conducted gene enrichment analysis. DEGs were enriched for a variety of biological categories, including neuronal function (e.g., neuropeptides), cytoskeletal organization, signaling (e.g., GPCRs), and extracellular matrix (**Fig. 5C-D, File S8-11**). To validate these effects in downstream GABA MNs of *unc-3(-)* mutants, we independently confirmed dysregulation of genes involved in vesicle sorting and trafficking (*unc-46/LAMP*)^12^, synapse remodeling (*oig-1*)^68^, and transcriptional regulation (*mab-9/TBX20*) using fluorescent reporters (**Fig. S8**). Altogether, these findings suggest that UNC-3 non-cell-autonomously impacts various biological processes in downstream GABA MNs.

### UNC-3 impacts neuropeptidergic signaling in downstream GABA MNs

Studies in invertebrates^69–76^ and vertebrates^77–81^ have highlighted neuropeptides as important modulators of locomotion, with implications for MN disease^82–84^. Given that host and downstream GABA MNs express cognate neuropeptide-receptor pairs^33,85^, and that UNC-3 activates neuropeptide gene expression in host MNs^4,6,86^, we asked whether loss of *unc-3* in host MNs affects neuropeptidergic gene expression in downstream GABA MNs.

Indeed, we uncovered significant dysregulation of neuropeptide and neuropeptide receptor genes both in host (79 genes) and downstream (15 genes) MNs of *unc-3(-)* animals (**Fig. S9A-B**). For example, validation using an endogenous fluorescent reporter for the neuropeptide *nlp-11* showed increased expression in both MN populations of *unc-3(-)* animals (**Fig. S9C**). Among the nine neuropeptide receptor genes that were significantly dysregulated in GABA MNs, two were upregulated: the FMRFamide receptor *frpr-5* and the DroMyoSuppressin G-Protein Coupled Receptor (GPCR) *dmsr-6* (**Fig. S9B**), both of which are implicated in control of locomotive behaviors^87–90^. Interestingly, *frpr-5* is dysregulated in both host and downstream MNs of *unc-3(-)* mutants, whereas *dmsr-6* was only upregulated in downstream MNs (**Fig. S9B**), a finding we validated using an endogenous fluorescent reporter for *dmsr-6* (**Fig. 5E-F**).

We next evaluated whether the *dmsr-6* upregulation in downstream MNs can be rescued by reintroducing UNC-3 specifically in host MNs of *unc-3(-)* animals and found this to be the case (**Fig. 5E-F**), supporting the notion that UNC-3 acts non-cell autonomously to suppress neuropeptide receptor (*dmsr-6*) expression in downstream MNs.

Taken together, these results suggest a broad role for the terminal selector UNC-3 in controlling neuropeptidergic signaling within the *C. elegans* motor circuit via cell and non-cell autonomous mechanisms in host and downstream GABA MNs, respectively.

### The UNC-3 effects on downstream GABA MNs depend on cholinergic neurotransmission

How can the loss of *unc-3* in host MNs lead to gene expression changes in downstream MNs? One possibility is that UNC-3 functions early in the GABA MN lineage such that its loss significantly alters the identity of post-mitotic GABA MNs. This possibility seems unlikely however, as our analysis of single-cell transcriptional^65^ and proteomic^66^ atlases throughout embryogenesis failed to detect appreciable levels of *unc-3*/UNC-3 early in the DD or VD MN lineages (**Fig. S7A-B’’**).

Another possibility is that UNC-3 acts in host MNs to control cell-cell signaling pathways (e.g., synaptic signaling or secreted cues) essential for downstream GABA MN development or function. To address this possibility, we focused on *dmsr-6* as a test case because it is specifically upregulated in downstream MNs of *unc-3(-)* animals (**Fig. 5E-F**). Because host MNs provide cholinergic (excitatory) input to downstream GABA MNs and UNC-3 controls the expression of ACh biosynthesis genes in host MNs^4,5,8^, we first asked whether blocking cholinergic transmission alone is sufficient to induce upregulation of *dmsr-6* in downstream MNs. Indeed, loss of cholinergic transmission either globally using a temperature sensitive allele *(y266)* for *cha-1* (*ChAT*), or specifically in host MNs using a regulatory mutation *(e113*, **Fig. S10**)^4,36^ upstream of the conserved *unc-17* (*VAChT*)/*cha-1* (*ChAT*) gene locus increased *dmsr-6* expression in GABA MNs (**Fig. 5G**). We conclude that *dmsr-6* upregulation in downstream MNs depends on UNC-3-controlled cholinergic neurotransmission from host MNs.

Next, we asked whether additional cell-cell signaling mechanisms downstream of UNC-3 might contribute to *dmsr-6* upregulation in GABA MNs. Since neuropeptide gene expression is significantly disrupted in both host and GABA MNs in the absence of UNC-3 (**Fig. S9**), we next investigated whether neuropeptide release also affects *dmsr-6* expression. Homozygous animals for a null allele *(e928)* of *unc-31*/*CAPS*, a gene required for neuronal neuropeptide release by dense core vesicles^91^, showed no effect on *dmsr-6* expression in GABA MNs (**Fig. 5H**). Further, *unc-3(-); unc-31(e928)* double mutants did not suppress the effect seen in *unc-3(-)* single mutants (**Fig. 5H**). We conclude that UNC-3 influence on gene expression in downstream MNs depends on cholinergic input from host MNs and is independent of neuronal neuropeptide release.

### Loss of *unc-3* triggers a CEBP-1-mediated transcriptional response in downstream MNs

The observation that cholinergic (host) MN input is sufficient to regulate *dmsr-6* expression in downstream GABA MNs suggests the involvement of neuronal activity-dependent transcriptional programs^92^. However, we detected no changes in the expression of established neuronal activity-dependent TFs (e.g., *crh-1/*CREB^93^, *fos-1/ATF2*^94^*, egl-43/MECOM*^94^, **File S2**) in downstream MNs of *unc-3(-)* animals. We next investigated several stress-induced TFs but found no dysregulation at the transcript level (e.g., *daf-16/FOXO4*^95^*, hsf-1/HSF1-2*^96^*, ets-5/FEV/Pet1*^97^, **File S2**).

To identify in an unbiased manner candidate TFs that mediate non-cell-autonomous transcriptional changes in downstream MNs of *unc-3(-)* animals, we used the recently published *Cel*Est gene regulatory networks (GRNs) inference algorithm^98^ (see Methods). As a control, we first tested the approach on host MNs. *Cel*Est correctly identified UNC-3 as the dominant driver of transcriptional changes in these cells (**Fig. 6A**). When applied to downstream MNs, the same unbiased analysis identified the conserved CCAAT/enhancer-binding protein (C/EBP) family bZIP TF CEBP-1 as the top candidate driver (**Fig. 6A**). Consistent with this prediction, *cebp-1* expression was sharply upregulated in neighboring MNs of *unc-3(-)* mutants (**File S2**). Using an endogenous translational reporter, we confirmed CEBP-1 upregulation in both DD and VD GABA MNs in *unc-3* mutants and observed a similar increase in its expression using the *unc-17(e113)* regulatory allele (**Fig. 6B-D**), indicating that reduced cholinergic transmission itself is sufficient to induce CEBP-1 in downstream MNs. Integration of our *unc-3(-)* scRNA-seq dataset with a published CEBP-1 ChIP-seq dataset^99^ revealed that ∼20% (40/212) of CEBP-1 genomic targets were dysregulated in GABA MNs, all of which were upregulated (**Fig. 6E-F**). Given that *cebp-1* (C/EBP) plays well-established roles in neuronal damage and regeneration in both *C. elegans*^100^ and vertebrates^101–103^, these findings support the notion that loss of *unc-3* or disruption of cholinergic transmission from host MNs triggers a CEBP-1-mediated neuronal injury/repair response in downstream GABA MNs.

**Figure 6.**
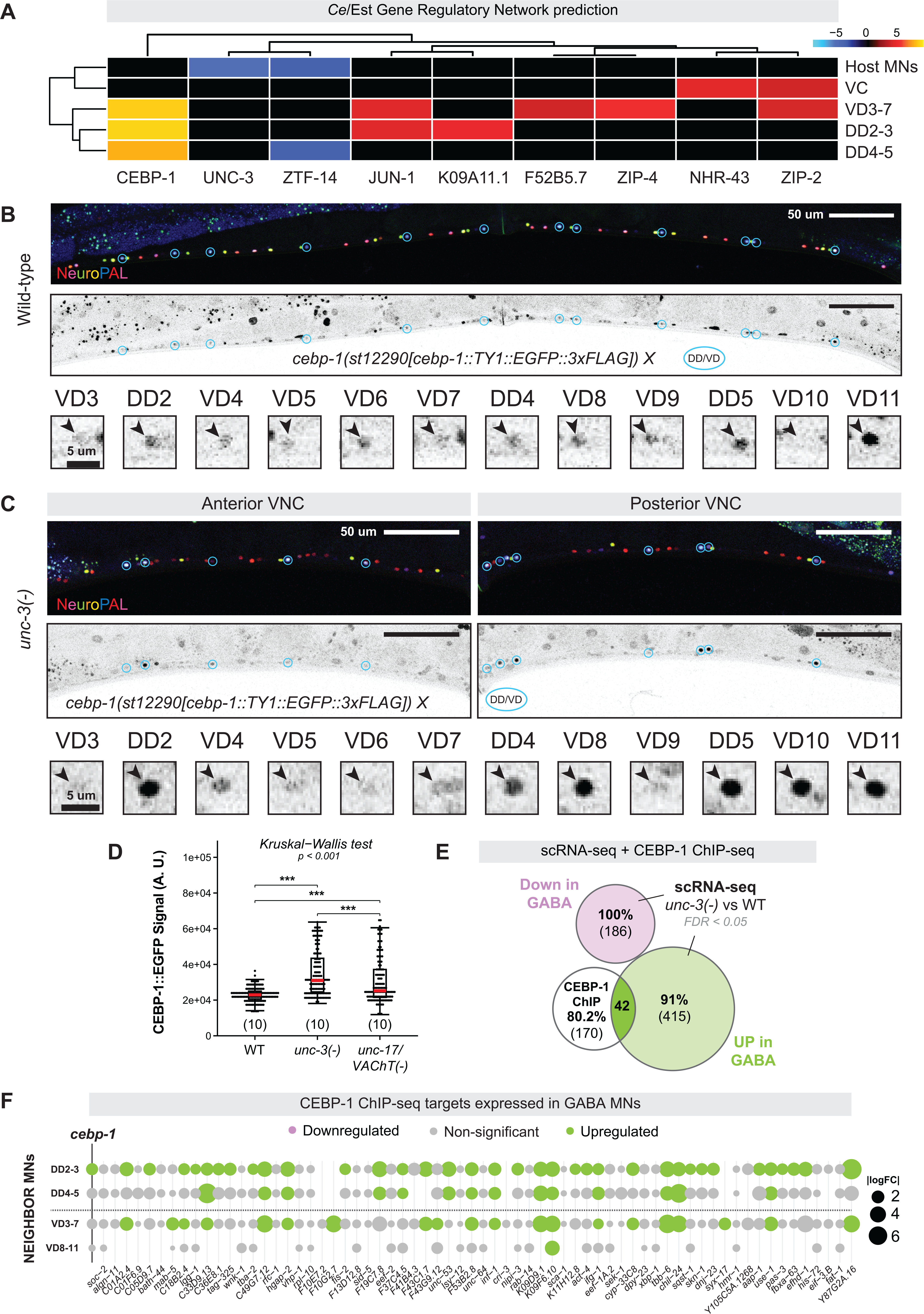
Loss of *unc-3* in host MNs triggers a pro-regenerative response in downstream MNs. **(A)** Heatmap showing *Cel*Est^98^ estimated TF activity (columns) driving differential expression between WT and *unc-3(-)* genotypes for host and downstream MNs (rows). Positive values (red-yellow) indicate increased transcription factor activity, whereas negative values (blue) indicate decreased activity. **(B-C)** Representative micrographs (top) of endogenous CEBP-1::EGFP expression in WT (A) and *unc-3(-)* (B) animals, with insets (bottom) of DD and VD MNs. **(D)** Fluorescence intensity quantifications of CEBP-1::EGFP in GABA MNs of WT, *unc-3(-)*, and *unc-17(e113)* animals. Quantifications were performed at young adult stage. N= 10 animals per genotype. Groups were statistically compared globally via a Kruskal-Wallis test, with *p*-values provided for significant and relevant comparisons; p>.05 = Not significant, ns; p<.05 = *; p<.01 = **; p<.001 = ***. Box-plot elements: thick horizontal line (red) = median; box = 25th to 75th percentiles (interquartile range); whiskers extend to the furthest data point within 1.5 × IQR from the quartiles (or the min/max if all points lie within this range); individual data points are overlaid as black dots. **(E)** Venn diagram depicting the integration of DEGs from scRNA-seq (this study) with CEBP-1 ChIP-seq data^99^. **(F)** Dot plot of CEBP-1 ChIP-seq targets^99^ that are expressed in GABA/downstreams MNs. Dot sizes convey absolute value of the log fold change (|logFC|) in gene expression, and dot color conveys regulation (upregulated in mutant = green; downregulated in mutant = pink). We note that all CEBP-1 target gene transcripts that were dysregulated in *unc-3(-)* animals were upregulated (green).

### Loss of *unc-3* in host MNs leads to axon pathfinding defects in downstream MNs

The significant transcriptomic effects we observed in downstream GABA MNs prompted assessment of their morphology in *unc-3(-)* animals. Using a GABA MN-specific fluorescent reporter *(juIs223[ttr-39p::mCherry]*), we identified four distinct phenotypes.

First, we observed axon guidance defects. In WT animals, GABA MN commissures typically project on the right side of the animal to reach the dorsal nerve cord^34,35^. In *unc-3(-)* animals however, 20% of GABA MN axons were located on the wrong (left) side (i.e., ‘reversed’) at both early larval (L1) and young adult stages (**Fig. 7A, S11A-B**). These reversals occurred without changes in the total number of commissures, suggesting an axon guidance phenotype (**Fig. 7A, S11C, E**). Second, whereas WT GABA MN axons bundle tightly together in the VNC, their axons display splitting or ‘defasciculation’ phenotypes in the *unc-3(-)* animals (**Fig. 7B**), reminiscent of cell adhesion defects^36,104,105^. Third, we observed increased GABA MN axonal gaps in the ventral nerve cord (**Fig. 7B-C**). Last, GABA MNs consistently exhibited axonal misrouting and ectopic branching around the vulva in *unc-3(-)* animals (**Fig. 7D-E**).

**Figure 7.**
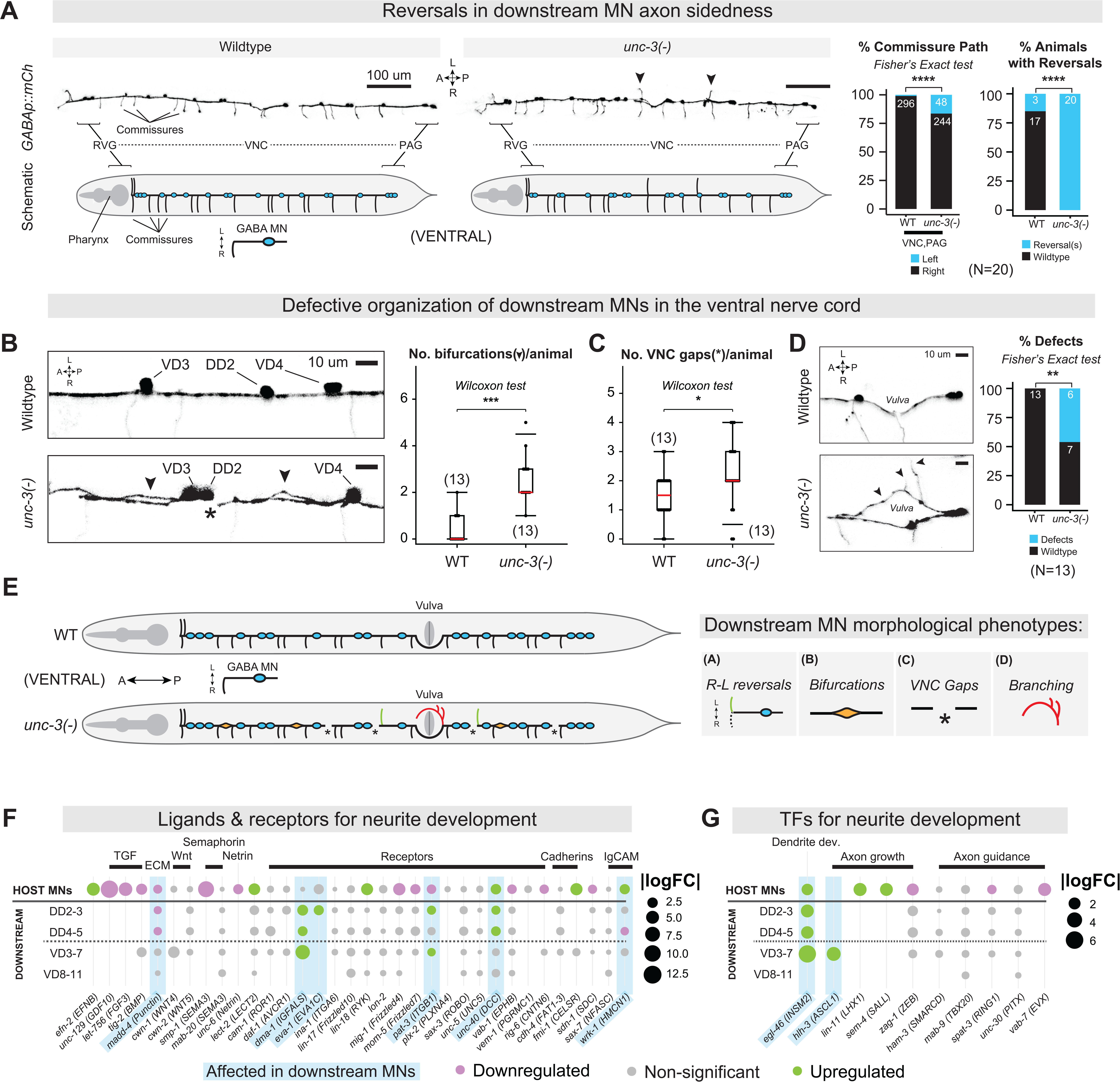
UNC-3/EBF impacts downstream cell morphology. **(A)** Left: Representative fluorescence micrographs of *ttr-39p::mCherry* expression in GABA VNC MNs seen ventrally (top) with schematic representations (bottom). Right: Percentage of right- vs. left-sided axonal/commissural paths and percentage of scored animals with this defect in wild-type vs. *unc-3(-)* backgrounds. Compared via Fisher’s exact test, p>.05 = Not significant, ns; p<.05 = *; p<.01 = **; p<.001 = ***; p<.00001 = ****. **(B-C)** Representative micrographs of GABA MN morphology in WT vs. *unc-3(-)* animals. Mutants display significantly more splits in the nerve fascicle (arrowheads) and more gaps in the nerve cords (asterisks, C) as visualized using the GABA MN-specific reporter. Quantifications were performed at young adult stage. Number of animals (N) is listed on each plot. Wilcoxon rank-sum test was selected based on non-normal data; p<.05 = *; p<.01 = **; p<.001 = ***. Box-plot elements: thick horizontal line (red) = median; box = 25th to 75th percentiles (interquartile range); whiskers extend to the furthest data point within 1.5 × IQR from the quartiles (or the min/max if all points lie within this range); individual data points are overlaid as black dots. **(D)** Left: Representative micrographs of ectopic branching near the vulva in *unc-3(-)* compared to WT. Right: Stacked bar plot showing percent of animals with defects. N = 13 animals. Compared via Fisher’s exact test, p>.5 = Not significant, ns; p<.05 = *; p<.01 = **; p<.001 = ***; p<.00001 = ****. **(E)** Schematic summary of morphological defects in GABA MNs of *unc-3(-)* animals assessed in A-D. **(F-G)** Dot plots showing DEGs encoding ligands and receptors (F) or TFs (G) associated with neurite (i.e., axon, dendrite) development. Gene list was derived from a recent review^106^. Dot sizes convey absolute value of the log fold change (|logFC|) in gene expression, and dot color conveys regulation (upregulated in mutant = green; downregulated in mutant = pink).

What is the molecular foundation for these axon pathfinding defects? To address this question, we cross-referenced our scRNA-seq dataset with a comprehensive list of *in vivo* validated genes that control neurite development (reviewed here^106^). We found dysregulation of neurite development genes in GABA MNs of *unc-3(-)* animals (**Fig. 7F**), including integrin (*pat-3/ITGB1*) and the Netrin receptor (*unc-40/DCC*). We also observed dysregulation of genes that mediate axonal (*eva-1/EVA1C, wrk-1/HMCN1*) and dendritic (*dma-1/IGFALS*) guidance. Finally, genes encoding TFs that regulate neurite development^106^ and regeneration^100,107,108^ (*egl-46/INSM2, hlh-3/ASCL1*) were also dysregulated in GABA MNs (**Fig. 7G**), revealing potential molecular mediators of the morphological phenotypes identified in *unc-3(-)* animals. Interestingly, we also found dysregulation of neurite development genes (e.g., *unc-6/Netrin*, *unc-40*/DCC, *wrk-1/HMCN1*) in host MNs (**Fig. 7F-G**), likely explaining the previously described axonal defects of *unc-3(-)* host MNs (**Fig. 1D**)^4,58,67^. Altogether, both transcriptomic and morphological analyses support a non-cell-autonomous role for UNC-3 in coordinating axonal/neurite development in downstream GABA MNs.

### UNC-3 influences downstream MN connectivity at the molecular and anatomical level

To assess whether the connectivity of downstream GABA MNs is affected in *unc-3(-)* animals, we first cross-referenced our scRNA-seq dataset with two comprehensive lists of *C. elegans* synaptogenesis molecules^109,110^. We found that both host and downstream cells exhibited changes in pre- and postsynaptic genes, which govern neuronal output and input^111^, respectively, (**Fig. 8A-B, S12**). In the GABA MN circuitry, most outputs are onto body wall muscles (BWMs) and most inputs come from cholinergic host MNs^34,35^ (**Fig. S13**). Importantly, GABA MNs form almost no gap (i.e., electrical) junctions with host MNs (**Fig. S14A**)^51,54^. Consistently, genes encoding innexins or other gap junction subunits were not affected in GABA MNs in the absence of *unc-3* (**Fig. S14B**).

**Figure 8.**
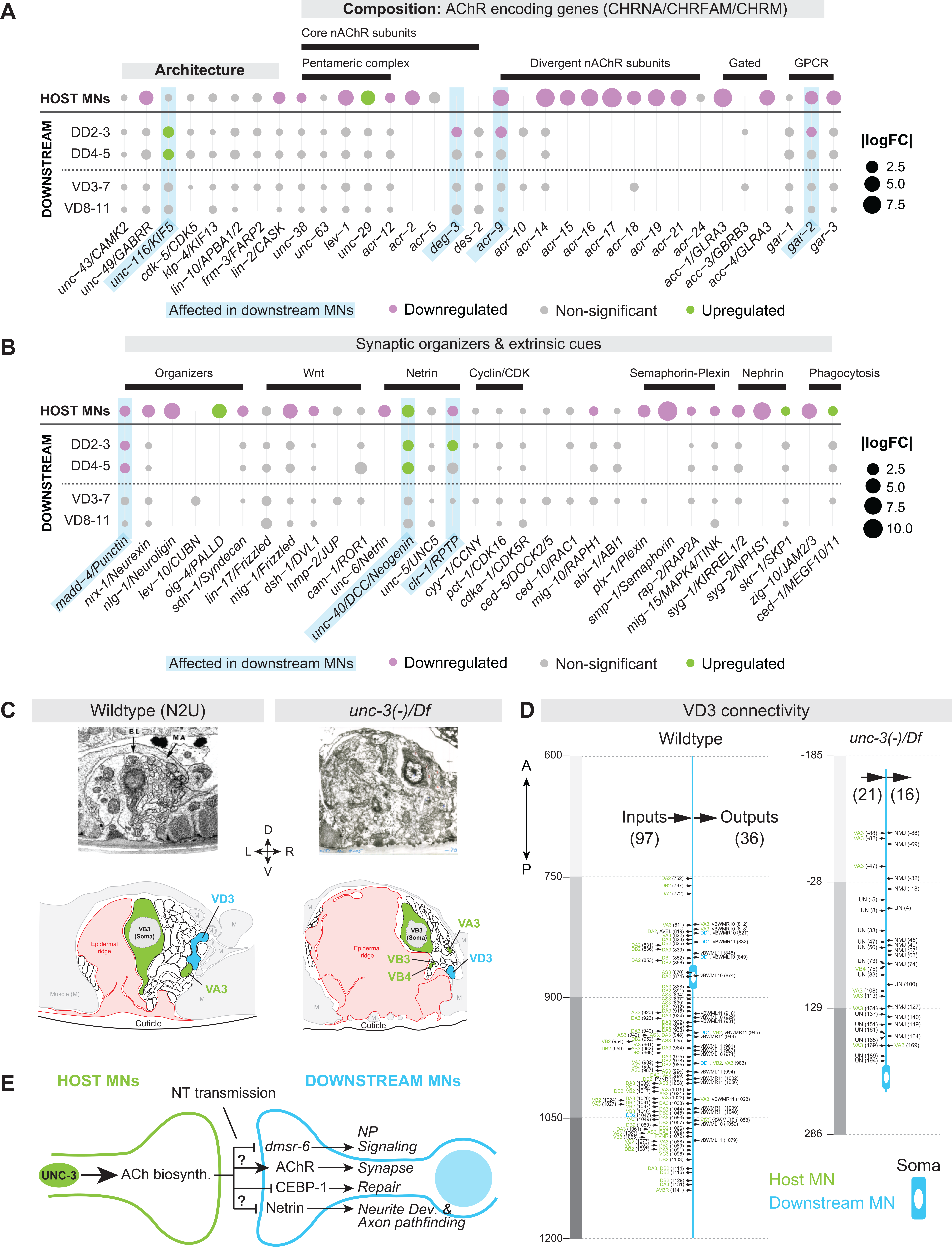
UNC-3/EBF impacts downstream MN connectivity. **(A-B)** Dot plots showing DEGs encoding postsynaptic components (A) and synaptic organizers and extrinsic cues for synapse development (B). Dot size conveys absolute value of the log fold change (|FC|) in gene expression, and dot color conveys regulation (upregulated = green; downregulated = pink). **(C)** Representative electron microscopy (EM) micrographs of cross-sections of a WT (N2U) and an *unc-3(-)/Df* animal (top) with schematic representations (bottom). **(D)** Synaptic inputs (left of axon/soma) and outputs (right of axon/soma) of the VD3 GABA MN in the WT and *unc-3(-)/Df* backgrounds. **(E)** Summary of scRNA-seq and genetic analyses, showing a role for cholinergic input in the non-cell-autonomous dysregulation of neuropeptide signaling (e.g., *dmsr-6*), neuronal injury/repair (CEBP-1 and effectors), synaptogenesis (e.g., AChR genes *acr-9, deg-3*), and axon pathfinding/neurite development (e.g., *unc-40/DCC, clr-1/RPTP*) in downstream MNs. Question marks indicate processes/pathways (AChR and Netrin genes) that are non-cell-autonomously impacted by UNC-3 but were not specifically assessed for loss of cholinergic signaling alone.

In DD MNs, expression of two genes implicated in synaptic output^112,113^ - *snt-6/SYT7* (Synaptotagmin) and *sng-1/SYNGR2/4* (Synaptogyrin) - was reduced (**Fig. S12A, File S2**). Expression of genes that mediate synaptic inputs was also disrupted in DDs (**Fig. 8A**), including a reduction in ACh receptor (AChR) genes, which mediate host MN signaling to downstream MNs. Specifically, three AChR genes, *deg-3/CHRNA7-2, acr-9*/*CHRFAM7A/CHRNA7* and *gar-2*/*CHRM3*, were downregulated (**Fig. 8A**). Interestingly, expression of transcripts for the five subunits of the core pentameric AChR/CHRNA complex (*acr-12, unc-38, unc-63, unc-29, lev-1*)^37^ was unaffected in downstream cells (**Fig. 8A**), despite the reported mislocalization of their corresponding proteins in DD MNs of *unc-3(-)* mutants^37,114^.

Given these changes in synaptic gene expression, we next evaluated the expression of 31 conserved genes involved in extrinsic pathways known to guide synapse formation in *C. elegans*^109^. Three - *unc-40/DCC, clr-1/RPTP,* and *madd-4/Punctin* - were non-cell-autonomously affected in DD MNs (**Fig. 8B**). The first two genes, *unc-40/DCC* and *clr-1/RPTP*, were upregulated in the mutant and encode receptors that together with their ligand UNC-6/Netrin, form a Netrin signaling pathway critical for synaptic partner recognition in other neurons in *C. elegans*^115^, suggesting a similar mechanism may mediate downstream MN synaptic connectivity. The third gene, *madd-4/Punctin*, was downregulated and encodes a secreted synaptic organizer from MNs that induces GABA receptor clustering in postsynaptic muscle cells^5,116,117^. Together, these results suggest that *unc-3* non-cell-autonomously impacts molecular features that determine synaptic input and output of downstream GABA MNs.

To complement this molecular analysis, we analyzed electron microscopy (EM) data (see Methods). Due to the axon pathfinding defects, we were unable to identify with confidence the DD neurons in the *unc-3(-)* EM dataset. Because a previous study found pre- and post-synaptic protein localization defects in VD neurons of *unc-3(-)* animals^36^, we reconstructed the synaptic connectivity of VD3 and VD4 from WT and *unc-3* mutant animals carrying a strong LOF allele (*e151*) over a chromosomal deficiency (*MnH205),* (hereafter *unc-3(-)/Df*) (**Fig. 1A, 8D**, **S15A**). The EM analysis revealed a marked reduction in inputs and outputs of VD3 and VD4, suggesting impaired integration into motor circuitry (**Fig. 8D, S15A**).

Reduced inputs onto the GABA VD MNs suggested that their presynaptic partners – the cholinergic host MNs - also have connectivity defects. Indeed, our analysis of six host cholinergic MNs (AS2, DA2, DB3, VA3, VB3, VB4) in the *unc-3(-)/Df* animal showed sharp reductions in their inputs and outputs (**Fig. S15B-G**), in addition to axon pathfinding defects (**Fig. S15H**). Altogether, the integration of EM and scRNA-seq datasets revealed that UNC-3 impacts the connectivity of downstream GABA MNs both at the anatomical and molecular level.

## DISCUSSION

Decades of research have emphasized the cell-autonomous roles of transcription factors (TFs) in nervous system development, but their impact on downstream, synaptically connected neurons remains poorly understood. Here, we employed scRNA-seq to systematically identify the cell and non-cell autonomous roles of the conserved terminal selector UNC-3/EBF1-4 in *C. elegans* ventral nerve cord motor neurons (MNs). The unbiased nature of scRNA-seq revealed a dual cell-autonomous role for UNC-3 in host MNs as both a transcriptional activator and repressor, as well as striking non-cell autonomous effects on the development of downstream GABA MNs. Since mutations in the *unc-3* human ortholog *EBF3* cause a severe neurodevelopmental syndrome (HADDS) characterized by muscle hypotonia and motor developmental delay^39–45^, the identification of new UNC-3 roles in host and downstream MNs may provide a conceptual framework for understanding the syndrome’s pathogenesis.

### UNC-3/EBF MN class-specific effects offer insight into neuronal diversification

Every nervous system is composed of individual neuron types which are often characterized by heterogeneity. That is, cells of a given neuron type share common features such as the use of the same neurotransmitter (NT) but can be further subdivided into distinct subtypes based on anatomical, functional, and molecular criteria. Terminal selectors tend to be expressed in all subtypes of a given neuron type^18^. For example, in mice, the terminal selectors *Pet-1* and *Nurr1* are expressed in all subtypes of brainstem serotonergic neurons^118^ and midbrain dopamine neurons^119^, respectively, but their precise functions in each subtype remain poorly understood. This highlights a conceptual problem broadly applicable to most neuron types: A common TF (e.g., terminal selector) specifies a group of neurons with shared features, but additional gene regulatory mechanisms must exist to diversify those neurons into subtypes. To date, our understanding of how terminal selectors control neuronal diversification processes, such as the identity of individual neuronal subtypes, remains limited due to the employment of either biased (candidate gene) or bulk RNA-seq approaches for the identification of TF target genes in post-mitotic neurons^120–125^. Using single-cell RNA-seq, we provide here a systematic characterization of terminal selector (*unc-3*) loss in all five classes (subtypes) of cholinergic MNs. Without *unc-3*, four of the five host MN classes (DA, DB, VA, VB), which control backward and forward locomotion^35^, collapsed into three molecularly distinct mutant populations (**Fig. 4G**). In contrast, the AS MN class, which has a functionally distinct role in movement coordination (i.e., speed, body curvature)^55,126^, showed resilience despite *unc-3* loss, partially retaining its original molecular features. This outcome is likely due to additional regulatory mechanisms, such as TFs, that help safeguard AS identity by partially compensating for *unc-3* loss. The strongest candidate is UNC-55/NR2F, which is known to control AS identity^7^, followed by ELT-1/GATA and MAB-9/TBX20, which are both expressed in AS neurons (**Fig. 2I**). Altogether, the single-cell resolution of our RNA-seq approach revealed a non-uniform picture likely applicable to other terminal selectors. Although all five host MN classes (DA, DB, VA, VB, and AS) are cholinergic and located in the nerve cord, loss of the same terminal selector (*unc-3*) led to differential effects on class (subtype) identity, offering valuable insights into the transcriptional logic of neuronal diversification.

### The dual role of UNC-3/EBF in host MNs expands the regulatory scope of terminal selectors

The unbiased identification of UNC-3 target genes in host MNs revealed a dual regulatory role. Consistent with prior work^4,6–8^, UNC-3 is a major determinant of MN identity by acting as a transcriptional activator of > 900 genes. However, our analysis also points to a new major role in transcriptional repression. Among the > 1,100 repressed genes in host MNs (**Fig. 3A**), we identified alternate neurotransmitter (NT) identity components. In *unc-3* mutants, host MNs ectopically express GABAergic or glutamatergic features (**Fig. 4**), suggesting that UNC-3 repressive function reinforces host MN identity by preventing the adoption of alternate NT identity features. This finding is reminiscent of other TFs in the vertebrate nervous system^127,128^. In mammalian serotonergic neurons, the terminal selector *Pet-1* activates serotonin pathway genes (*Tph2*, *Slc6a4/SERT)* and inhibits alternate NT identities (dopaminergic or noradrenergic)^129–131^. In the developing cerebellum and spinal cord, *Ptf1a* drives GABAergic identity and suppresses glutamatergic identity^132,133^. However, it has remained unclear whether the repressor activity of terminal selectors is continuously needed or becomes superfluous in later stages of life. With inducible protein depletion assays, we found that the UNC-3 repressive function is continuously required to prevent alternate NT identities in mature cholinergic MNs (**Fig. 4H-L**).

The roughly equal numbers of activated (929) and repressed (1,121) target genes in host MNs suggest that gene repression is a major function of UNC-3 that likely extends to other *C. elegans* neuron types that express *unc-3*, such as the head ASI chemosensory neurons^134^. But how can UNC-3 activate and repress distinct sets of genes? Prior studies found that terminal selectors can suppress inappropriate neuronal identities in their host cells via TF competition mechanisms^135,136^. Alternatively, they could prevent the expression of other terminal selectors. However, this does not appear to be the case for UNC-3, as we found no ectopic expression of terminal selectors of GABA (*unc-30*) or glutamatergic (*che-1, unc-86, die-1*) NT identities in *unc-3(-)* host MNs, nor evidence of UNC-3 binding in their *cis*-regulatory regions (**Fig. S3A-B**). Instead, we found that UNC-3 binds directly to 36% of its repressed target genes, suggesting a direct transcriptional repressor role. Whereas prior work established that the cognate COE binding motif (TCCCNNGGA) of UNC-3 is required for gene activation in host MNs^4,5,8^, its newly uncovered repressor function differs from its activator role distinct in two distinct ways: it is largely independent of its cognate (canonical) motif and exhibits promoter bias (**Fig. 3 E-F**). Future mechanistic studies are needed to determine whether UNC-3 cooperates with other TFs to bind the non-cognate motifs identified here (**File S6**) to repress its targets in host cells. Last, UNC-3 promoter bias in the *cis*-regulatory sequences of repressed targets may reflect an underlying position-dependent mechanism, as was recently described for human TFs^137^.

We emphasize that the relatively low percentage (36%) of repressed genes bound by UNC-3 suggests that UNC-3 predominantly acts indirectly to repress genes in host MNs. Consistent with this idea, we previously found that UNC-3 antagonizes the Hox protein LIN-39 (Scr/Dfd/Hox4-5), preventing it from activating inappropriate terminal identity genes in host MNs^9^. Altogether, our work offers a blueprint for future single-cell studies on terminal selectors, aiming to resolve the mechanisms through which these conserved and clinically relevant TFs sculpt neuron identity programs.

### Non-cell-autonomous roles of UNC-3/EBF in shaping downstream MN development

A major challenge in understanding animal development has been the identification of both cell and non-cell autonomous mechanisms through which TFs control cell type identity. These specific effects could be especially important for the development and maturation of post-mitotic neurons in the emerging nervous system. Here, we show that UNC-3/EBF1-4 non-cell-autonomously affects the transcriptomes of downstream synaptically connected GABA (DD and VD) MNs in the *C. elegans* ventral nerve cord, indirectly modulating transcription of genes involved in key neuronal processes such as neuropeptidergic signaling and neurite development. These molecular results, supported by axon pathfinding (**Fig. 7**) and synapse ultrastructural (**Fig. 8**) analyses, expand the conventional view of terminal selectors as solely intrinsic regulators of host neuron identity^2,16–19^. Further, our findings establish the *C. elegans* motor circuit as a powerful model for studying causal pathways linking TF activity in host cells to its impact on downstream cells.

Given that *unc-3* is not expressed in the lineage of GABA (DD and VD) MNs (**Fig. S7**), we surmise that cell-cell signaling from host to downstream MNs is disrupted in *unc-3* mutants, leading to the observed molecular and morphological effects in DD and VD cells. In principle, such cell-cell signaling can take many forms, ranging from synaptic signaling to secreted molecules and cell surface proteins. Here, we find that cholinergic neurotransmission from host to downstream MNs is sufficient to induce transcriptional changes in GABA MNs (**Fig. 8E**), consistent with prior work showing that cholinergic host MN input drives synaptic structural changes in DD^138^ and VD MNs^36^. Although cholinergic neurotransmission is involved, we detected no changes in expression of established neuronal activity-dependent TFs (e.g., *crh-1/*CREB^93^, *fos-1/ATF2*^94^*, egl-43/MECOM*^94^) in GABA MNs of *unc-3(-)* animals. Instead, we found increased expression of the pro-regenerative bZIP TF CEBP-1^100,107^ in *unc-3* and *unc-17/VAChT* mutants, supporting the idea that disruption of cholinergic neurotransmission in host MNs, which is a consequence of *unc-3* loss, triggers a CEBP-1-mediated neuronal injury/repair response in downstream GABA MNs (**Fig. 8E**).

Notably, we also observed dysregulation of genes encoding major components of the deeply conserved UNC-6/Netrin signaling pathway in both host and downstream MNs of *unc-3(-)* animals (**Fig. 8B**). Netrin signaling regulates a variety of neuronal processes from nematodes to mammals, including axon guidance^139,140^, neurite morphology^141,142^, and synapse assembly^115,143,144^, all of which appear defective in downstream MNs of *unc-3(-)* animals. In addition, UNC-3 is known to directly activate expression of the presynaptic cell-adhesion molecule *nrx-1*/Neurexin in host MNs; loss of UNC-3 strongly reduces NRX-1 levels leading to failure of dendritic spine formation and postsynaptic AChR receptor clustering in downstream GABA MNs^37^. We surmise that these two conserved non-cell-autonomous signaling pathways – UNC-6/Netrin and NRX-1/Neurexin – which are controlled by UNC-3 in host MNs, likely act in parallel to inform morphological and synaptic development in downstream GABA MNs. Interestingly, a similar logic has been described for the terminal selector UNC-42 (PROP1), which cell-autonomously controls host neuron identities and circuit assembly while non-cell autonomously promoting axonal outgrowth in neighbors, potentially via its control of UNC-6/Netrin^145,146^. Hence, the non-cell-autonomous effects of UNC-3 may reflect a broader role for terminal selectors in deploying conserved guidance and adhesion cues to coordinate neuronal circuit development.

### Limitations

Our study provides a detailed analysis of how the TF UNC-3/EBF affects molecular and cellular processes in both host and downstream MNs of the *C. elegans* ventral nerve cord, offering a rare glimpse into how TF activity in host cells affects the development of downstream cells. However, we do not explore the potential impact of UNC-3 on non-neuronal cells, such as muscle cells, which also receive cholinergic input from host MNs.

Consistent with prior work^67,146,147^, we found that *unc-3* loss impacts the organization of the ventral nerve cord (**Fig. 7**), which could secondarily alter the transcriptomes of downstream GABA MNs. While we cannot rule out that nerve cord disorganization contributes to the observed downstream cell defects, the minimal transcriptomic changes in the VC class of neurons, which are also located in the nerve cord, suggests that the non-cell-autonomous effects of UNC-3 are specific to GABA MNs.

## Conclusions

By uncovering regulatory roles for a conserved terminal selector gene, this work advances our understanding of how TFs control the development and function of individual neuron types, shifting the field’s perspective from a “host cell-centric” framework to one that incorporates unexpected non–cell-autonomous roles in neighboring neurons. Together, our findings offer a conceptual framework for understanding complex TF functions in neural development and circuit assembly, as well as the pathogenesis of the *EBF3* neurodevelopmental syndrome (HADDS).

## ACKNOWLEDGEMENTS

We thank the *Caenorhabditis* Genetics Center (CGC), which is funded by the National Institutes of Health (NIH) Office of Research Infrastructure Programs (P40 OD010440), for providing many of the strains used in this study. We thank members of the Kratsios lab (Filipe Marques, Konstantinos Tsioras, Ian Weigle), Oliver Hobert and Michael Francis for providing feedback on the manuscript. We also thank the Vanderbilt University Medical Center (VUMC) Flow Cytometry Shared Resource, supported by Ingram Cancer Center (P30 CA68485), DDRC (DK058404), VANTAGE Core Facility, supported by CTSA (5UL1 RR024975-03), Ingram Cancer Center (P30 CA68485), Vision Center (P30 EY08126), NIH/NCRR (G20 RR030956) for scRNA-Seq, and the Vanderbilt Cell Imaging Shared Resource (NIH CA68485, DL20593, DK58404, DK59637, EY08126). This work was funded by NIH grants to J.J.S. (K99 NS140549-01), P.K (R21 NS108505, R01 NS118078, R01 NS116365), D.H (OD R24 010943), and D.M.M. (R01NS113559, R01NS100547 and R01NS106951).

## AUTHOR CONTRIBUTIONS

J.J.S., S.R.T., D.M.M., and P.K. were responsible for conceptualization; J.J.S., S.R.T, G.K., and D.H.H. were responsible for investigation; J.J.S., S.R.T., G.K., and D.H.H. were responsible for formal analysis. J.J.S., D.M.M, and P.K. were responsible for resources and funding acquisition. J.J.S. and P.K. were responsible for writing the original draft. D.H.H., D.M.M., and P.K. were responsible for supervision. All authors read, edited, and approved of the manuscript.

## METHODS

### Maintenance of C. elegans strains

All *C. elegans* strains were cultured at 20°C or 25°C on nematode growth media (NGM) plates seeded with standard *E. coli* OP50 as a food source. Embryonic, larval, and day 1 adult stage hermaphrodites were assessed as described in the main text, figures, and figure legends. All strains used or generated for this study are listed in the **Key resources table**.

### Preparation of day 1 adult stage C. elegans and dissociation

Worms were grown on 8P nutrient agar plates (150 mm) seeded with *E. coli* strain NA22. To obtain synchronized cultures of day 1 adult stage animals, adult hermaphrodites were treated with hypochlorite to isolate embryos, which were then incubated in M9 buffer at room temperature for 16 hours to hatch and arrest at L1 stage. L1 animals were then grown on NA22-seeded plates for 62-65 hours. The developmental age of each culture was determined by scoring vulval morphology in at least 90 worms. The *lin-39p::RFP* strain showed variability in age even after synchronization. Thus, young adults were further isolated by size exclusion using a 35 µm nylon mesh. Worms were washed off plates with M9 then placed on a 35 µm nylon mesh suspended over M9 containing NA22 bacteria for 25 minutes. Larval worms pass through the mesh, whereas adults were retained. Young adult worms remaining on the mesh were collected into M9 and centrifuged at 150 relative centrifugal force (rcf) for 2.5 minutes. Of the remaining *lin-39p::RFP* population, 81.5% of the animals were young adults and the remaining 18.5% were late L4 stage (approaching day 1 adult stage). Single-cell suspensions were obtained with some modifications to published protocols^148,149^. Worms were transferred to a 1.6 mL centrifuge tube and centrifuged at 16,000 rcf for 1 minute to form 250 uL pellets. Worm pellets were then treated with 500 µL of SDS-DTT solution (20 mM HEPES, 0.25% SDS, 200 mM DTT, 3% sucrose, pH 8.0) for 6 minutes.

Following SDS-DTT treatment, worms were washed five times (each wash: dilute with 1 mL egg buffer, centrifuge at 16,000 rcf for 30 seconds). Worms were then digested in cold-active protease (10 mg/mL, Protease from Bacillus licheniformis, Sigma-Aldrich P4860, with DNAse I 35 U/mL) in egg buffer for 35 minutes at 4°C. Cells were kept at 4°C from protease incubation through all subsequent procedures.

During the protease incubations, solutions were triturated by pipetting through a P1000 pipette tip for four sets of 80 repetitions. Dissociation efficiency was monitored at 5-minute intervals with a fluorescence dissecting scope. Digestions were stopped by adding 750 µL L-15 media supplemented with 10% fetal bovine serum (L-15-10). Cells were then centrifuged at 530 rcf for 5 minutes at 4°C. The pellet was resuspended in 1 mL L-15-10, and individual cells were separated from whole worms and debris by centrifuging at 100 rcf for 2 minutes at 4°C. The supernatant was filtered through a 35-micron strainer cap into a 5 mL collection tube (Falcon 352235). The pellet was resuspended a second time, centrifuged (100 rcf for 2 minutes at 4°C), and the resulting supernatant was added to the collection tube. For higher sorting efficiency, samples were diluted with an additional 1 mL of L-15-10.

### FACS isolation of C. elegans motor neurons for single-cell RNA-seq

The samples were prepared as three pairs isolated on different days from distinct cultures, with a WT (*lin-39p::RFP*) sample and *lin-39p::RFP; unc-3(n3435)* sample being prepared in parallel on each day. Single-cell suspensions of both genotypes were sorted simultaneously on two BD FACSAria™ III cell sorters faceted with 70-micron diameter nozzles. DAPI was added to the samples (final concentration of 1 µg/mL) to label dead /dying cells. Non-fluorescent N2 standards were used to set gates to exclude auto-fluorescent cells. To ensure capture of all targeted VNC MNs, we set FACS gates to encompass a wide range of fluorescent intensities. This likely contributed to the presence of unlabeled neuronal and non-neuronal cells in our, which were excluded from our analysis in this study. Cells were sorted for three hours under the “4-way Purity” mask into 1.5 mL microcentrifuge tubes containing 200 uL L-15-33 (L-15 medium with 33% fetal bovine serum) and subsequently centrifuged (1200 rcf for 12 minutes at 4°C). A hemocytometer was used to count fluorescent cells. Single-cell suspensions used for 10x Genomics single-cell sequencing ranged from 180-350 cells/µL. Three samples of WT and three samples of *unc-3(n3435)* were prepared for encapsulation using the 10X Genomics Chromium system. We estimate the time from harvesting worms off plates (dissociation + sorting = ∼4.5 hours) to encapsulation in the 10X Genomics Chromium Controller (concentrating + counting + encapsulation = 1-1.5 hours) to be ∼6 hours.

### Single-cell RNA sequencing

Each sample targeted 10,000 cells and was processed for single cell 3’ RNA sequencing using the 10X Chromium system (v3 chemistry). Libraries were prepared using P/N 1000075, 1000073, and 120262 following the manufacturer’s protocol. Illumina NovaSeq 6000 with 150 bp paired end reads was used for library sequencing. Base calling was performed with Real-Time Analysis software (RTA, version 2.4.11; Illumina).

### Downstream processing

FASTQ files were processed using the 10X Genomics Cell Ranger software (version 6.1.1) using a custom reference genome from Wormbase release WS273 that incorporated extended 3’ UTR regions for specific several genes^149^. This reference is accessible at GEO under accession number GSE234962. Cells containing droplets were identified via the Emptydrops function in the DropletUtils R package, with the ambient RNA profile derived from droplets containing fewer than 50 unique molecular identifiers (UMIs). Background RNA correction was performed with the SoupX R package^150^, applying a 25-UMIs cutoff for the background expression profile. The genes used for contamination estimation in each sample are detailed in **File S1**, column J. Adjusted counts were rounded to the nearest integer, and the resulting corrected gene-by-barcode matrices were carried forward for further processing. Quality control metrics were calculated for each dataset using the scater R package^151^, focusing on the proportion of UMIs attributed to mitochondrial genes (*nduo-1, nduo-2, nduo-3, nduo-4, nduo-5, nduo-6, ctc-1, ctc-2, ctc-3, ndfl-4, atp-6, ctb-1, MTCE.7* and *MTCE.33*). Droplets exceeding 20% mitochondrial UMI content were filtered out. Up to this stage, the six samples were processed separately. The datasets were then merged into a single cell_data_set object within the monocle3^152–155^ framework for dimensionality reduction and clustering. Dimensionality reduction on the integrated dataset used 75 principal components, followed by 2-dimensional UMAP embeddings with parameters of umap.min_dist = 0.3 and umap.n_neighbors = 75, which had previously demonstrated effective separation of most neuron classes^10^. All remaining parameters were default values. One WT sample – corresponding to the third sample preparation day – yielded low-quality cells and was therefore excluded from analysis.

### Motor neuron class and subclass identification in C. elegans

We identified clusters containing VNC MNs by expression of known marker genes, including *unc-3, bnc-1, unc-4, unc-129, vab-7, unc-25,* and *unc-47* (Fig. S1A, File S1). Clusters corresponding to GABAergic MNs (DD and VD) and to the VC neurons contained overlapping populations of cells from both genotypes. We could confidently annotate clusters corresponding to AS, DA, DB, VA and VB from WT, but these clusters did not contain any cells from *unc-3(n3435)* animals. We identified populations of cells we surmised to be cholinergic MNs in the *unc-3(n3435)* data by expression of neuronal markers *egl-3, egl-21* and *sbt-1* as well as the homeodomain TFs *ceh-20, unc-62,* and the Hox genes *ceh-13, lin-39* and *mab-5.* We subset the data to include only those populations corresponding to WT or *unc-3(n3435)* VNC MNs and reran dimensionality reduction (35 principal components), alignment with align_cds (from the batchelor R package, alignment_k = 10), and generated new UMAP plots (umap.min_dist = 0.2, umap.n_neighbors = 50). The subsequent sub-UMAP was also made of just the host MNs in both genotypes using 25 principal components, umap.min_dist = 0.2, umap.n_neighbors = 50). We note that while the *lin-39::RFP* reporter labeled most VNC MNs in WT and *unc-3(n3435)* animals (Fig. 1D-E), we did not sample each MN class across genotypes equally (File S1).

### Differential Expression in C. elegans motor neurons: wild-type vs. unc-3(n3435)

Differential expression analyses between WT and *unc-3(n3435)* MN classes were performed using pseudobulk methods. For each class, an aggregate “pseudobulk” expression value was calculated for every gene for each replicate sample (two WT and three *unc-3(n3435)* samples) using the aggregateAcrossCells function from the scater R package. In the case of neighbor MNs and AS neurons, each subclass (e.g., DD2_3, DD4_5, VC4_5) was identified in both genotypes and was therefore considered a separate class. In the case of host MNs, expression was aggregated across all DA, DB, VA, and VB neurons in the WT samples, and across all cells in Groups 1-3 in the *unc-3(n3435)* samples. The pseudobulk expression profiles were then used as input to edgeR for differential expression analysis. Each class was considered independently.

### DEG enrichment analysis

Gene set enrichment analysis was performed using WormCat2.0 (wormcat.com)^60^. For each host or downstream MN class, two separate gene lists were submitted: (1) genes significantly upregulated (activated) and (2) genes significantly downregulated (repressed), using the same statistical cutoffs applied for differential expression (False discover rate [FDR] < 0.05). WormCat2.0 was run with default settings using the “Fisher Exact Test” mode and the full ‘Whole genome v2’ annotation type. Categories from all three WormCat annotation levels (Category 1–3) were examined, and enriched terms with FDR < 0.05 were considered significant. Results were exported as CSV files (Files S8-11).

### Transcription factor activity estimation

We used the *Cel*EsT gene regulatory networks^98^ to estimate transcription factor (TF) activity. The *Cel*EsT GRNs contain TF and target data for 487 TFs, compiled from ChIP-Seq, DNA-binding motifs, and eY1H screens. For each MN subclass, we used the average log fold change between wild-type and *unc-3(n3435)* for all genes as input, along with the *Cel*EsT GRNs, to the decouple function in the decoupleR R package^156^. We used the multivariate linear model (“mlm”). We calculated Benjamini-Hochberg corrected *p*-values to control for multiple comparisons within each neuron. We retained only those transcription factors with adjusted *p*-values < 0.05. We also filtered results to retain TFs that were detected in the neuron of interest in at least one genotype.

### Microscopy

Larval and adult stage animals were anesthetized with sodium azide (NaN_3_, 100 mM) and mounted on a 4% agarose pad on imaging slides. Images were recorded with an automated fluorescence microscope (Zeiss, Axio Imager Z2). Z stacks (0.50-1.0 µm step size) were acquired using a Zeiss Axiocam 503 mono (ZEN Blue software, version 2.3.69.1000), a Zeiss LSM 880, or a Zeiss LSM 900. Representative images shown are max intensity z-projections. All image reconstruction was performed in Fiji (version 2.9.0/1.53t). Fluorescent overlays and cropping for figures were generated using Adobe Photoshop 2022 (23.2.2 Release). Images of strains containing the fluorescent Neuronal Polychromatic Atlas of Landmarks (NeuroPAL) transgene^157^ for cell identification were acquired on a Zeiss LSM900 confocal microscope with 0.5-1.0 µm step size.

### Fluorescence intensity quantifications

To quantify fluorescence intensity in individual VNC MNs, Z-stack images of the midbody or whole animal were acquired at 0.5-1.0 um z-step intervals. The same imaging parameters (e.g., exposure time, laser power, master gain, temperature) were applied across all samples within the same experiment. Image stacks were then processed, including background subtraction via sliding paraboloid at rolling ball radius 50.0 pixels, and signal was measure using FIJI.

### Identification of motor neuron classes and subclasses in C. elegans

Motor neuron classes and subclasses were identified based on the following criteria: (1) Anatomically invariant positioning of neuronal bodies in the retrovesicular ganglion (RVG), ventral nerve cord (VNC), and preanal ganglion (PAG); (2) class-specific birth order (i.e., embryonic versus post-embryonic emergence); (3) total cell numbers in each motor neuron class or subclass; (4) co-expression with the NeuroPAL transgene ^157^.

### Temporally controlled UNC-3 degradation

To deplete UNC-3 specifically in mature (L4-day 1 adult stage) animals, we used the auxin-inducible degradation system^64^ with the previously characterized KRA376 strain, which contains *ot837[unc-3::mNG::AID]* allele and ubiquitously driven TIR1 *ieSi57[eft-3::TIR1::mRuby]*^8^, crossed to *otIs564[unc-47^fosmid^::mCherry]*. L4 stage animals were grown at 20°C on NGM plates coated with 4 nM auxin (indole-acetic acid [IAA] dissolved in ethanol) or ethanol (negative control) for 24 hours before imaging. All plates were shielded from light.

### UNC-3 cDNA rescue in host MNs

A 558bp fragment of the *unc-3* promoter, which expresses in VNC host MNs, was fused to UNC-3 cDNA sequence followed by the *unc-54* 3’ UTR via PCR fusion. The *unc-3p::UNC-3(cDNA)::unc-54(3’UTR)* fusion fragment was microinjected into young adult WT (N2) hermaphrodite animals at 50 ng/uL with *ttx-3p::mCherry* as a co-injection marker (50 ng/uL). Animals harboring this construct were then crossed to the *unc- 3(n3435)* null mutant containing the NeuroPAL transgene and endogenous *dmsr- 6(syb4442[dmsr-6::SL2::GFP::G2B])* reporter for the rescue assay. As this *unc-3p::unc- 3* cDNA was maintained extra-chromosomally, cDNA(-) and cDNA(+) siblings were used to score rescue.

### Integrated scRNA-seq and ChIP-seq motif analysis

Peak calls from UNC-3 ChIP-seq narrowbed file^8^ were annotated to nearby genes with Hypergeometric Optimization of Motif EnRichement (HOMER) discovery and analysis tool using annotatePeaks.pl^158^. Annotated peaks were then filtered based on their association with either activated or repressed DEGs. *De novo* motif discovery was conducted on the filtered peak groups with HOMER using findMotifsGenome.pl, with a size of +/- 75bp from the center of each peak. Throughout this study, ChIP-seq tracks were generated using Integrated Genomics Viewer (IGV)^159,160^.

### Electron microscopical analysis

To assess neuronal morphology and synaptic connectivity, we performed reconstructions of WT and *unc-3* mutant animals in the anterior segment of the ventral nerve cord. The strains examined included the WT strain (N2U) and the *unc-3(e151)* variant balanced by the *mnDf5* deficiency. The *unc-3* mutant analyzed was a trans- heterozygous combination of the *unc-3(e151)* loss-of-function allele (depicted in Fig. 1A) with the chromosomal deficiency *mnDf5*, which fully deletes the *unc-3* locus. This balanced strain was created through a cross between *unc-3(e151)* and the SP266 line carrying mnDp1(X;V)/V; *mnDf5* X. Reconstructions were derived from serial-section electron micrographs outlined previously^51^. The analyzed regions spanned approximately 150 µm and encompassed around 1800 consecutive sections. Specifically, we focused on the VNC extending roughly from the AS1 to AS3 MNs. Images were captured and produced for every third section. We traced the full axonal and dendritic processes of host and downstream MNs whose somata resided within the reconstructed zone (Fig. 8, S15). Neuron identities were established based on distinctive synaptic patterns, morphological traits, and the sequential positioning of their cell bodies along the ventral cord^51^.

### Statistics and reproducibility

For continuous data (i.e., MN counts, fluorescent intensity of reporters, morphological assays), statistical tests were selected based on data distribution as follows: Normally distributed data (Shapiro-Wilk *p* > 0.05 in both groups) with equal variances (Levene’s test *p* > 0.05) = Student’s t-test; Normally distributed with unequal variances = Welch’s t-test; Non-normal data = Wilcoxon rank-sum test. Box-plot elements: thick horizontal line (red) = median; box = 25th to 75th percentiles (interquartile range); whiskers extend to the furthest data point within 1.5 × IQR from the quartiles (or the min/max if all points lie within this range); individual data points are overlaid as black dots. Fluorescent microscopy images next to each dot plot show a representative animal. The number of animals (N) is listed near each box in all plots. No data were excluded from analyses and investigators were not blinded to allocation during experiments and outcome assessment.

For statistical comparison of categorical data (i.e., UNC-3 ChIP-seq peak distribution in Fig. 3f, frequency of morphological defects in Fig. 7, S11), Fisher’s exact test was performed. The number of animals (N) is listed near each group in all plots. Where >2 categories were compared, post-hoc 2×2 Fisher’s exact tests with Bonferroni correction were performed. For all quantifications/statistical tests, p>.05 = not significant, ns; p<.05 = *; p<.01 = **; p<.001 = ***; p<.0001 = ****.

