## Supplementary material for "Intrinsic and non-cell autonomous roles for a neurodevelopmental syndrome-linked transcription factor": Figure S1

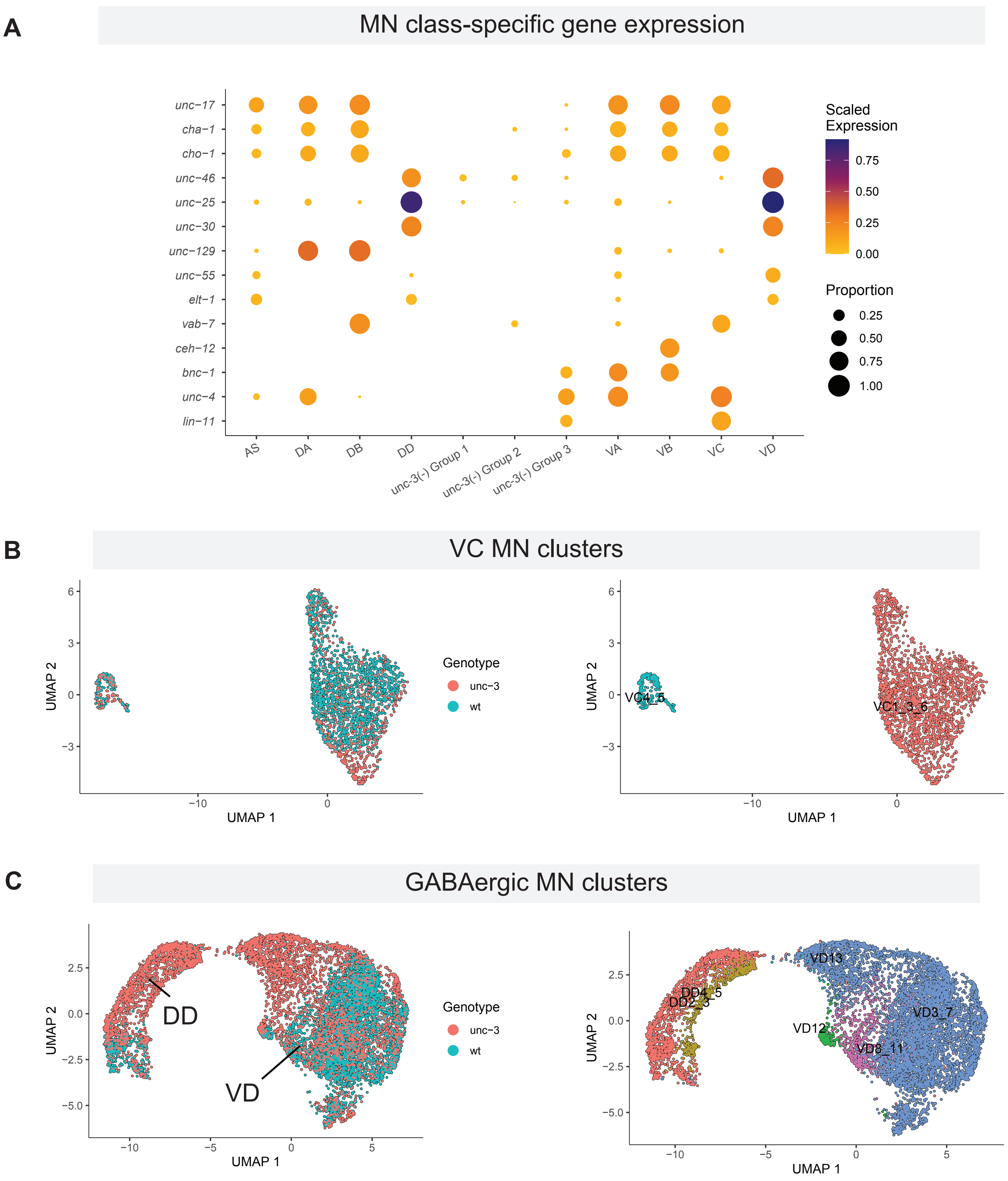


**Figure S1. Host MN class identity collapses in the absence of *unc-3* expression. (A)** Dotplot depicting expression of known MN class-specific genes in 5 WT host MN classes (DA, DB, VA, VB, AS), 3 downstream MN classes (DD, VD, VC) and in *unc-3(-)* mutant cells. Dot color indicates scaled gene expression and dot size indicates the proportion of cells in the cluster expressing the gene. **(B-C)** UMAP subsets depicting integration of WT and *unc-3(-)* VC (B) and GABA (C) MNs from scRNA-seq. Left: UMAP where colors indicate genotype from which each sequenced cell was derived (blue = WT; red = *unc-3(-)*). Right: UMAPs where colors indicate MN subclass identity.
