## Supplementary material for "Intrinsic and non-cell autonomous roles for a neurodevelopmental syndrome-linked transcription factor": Figure S2

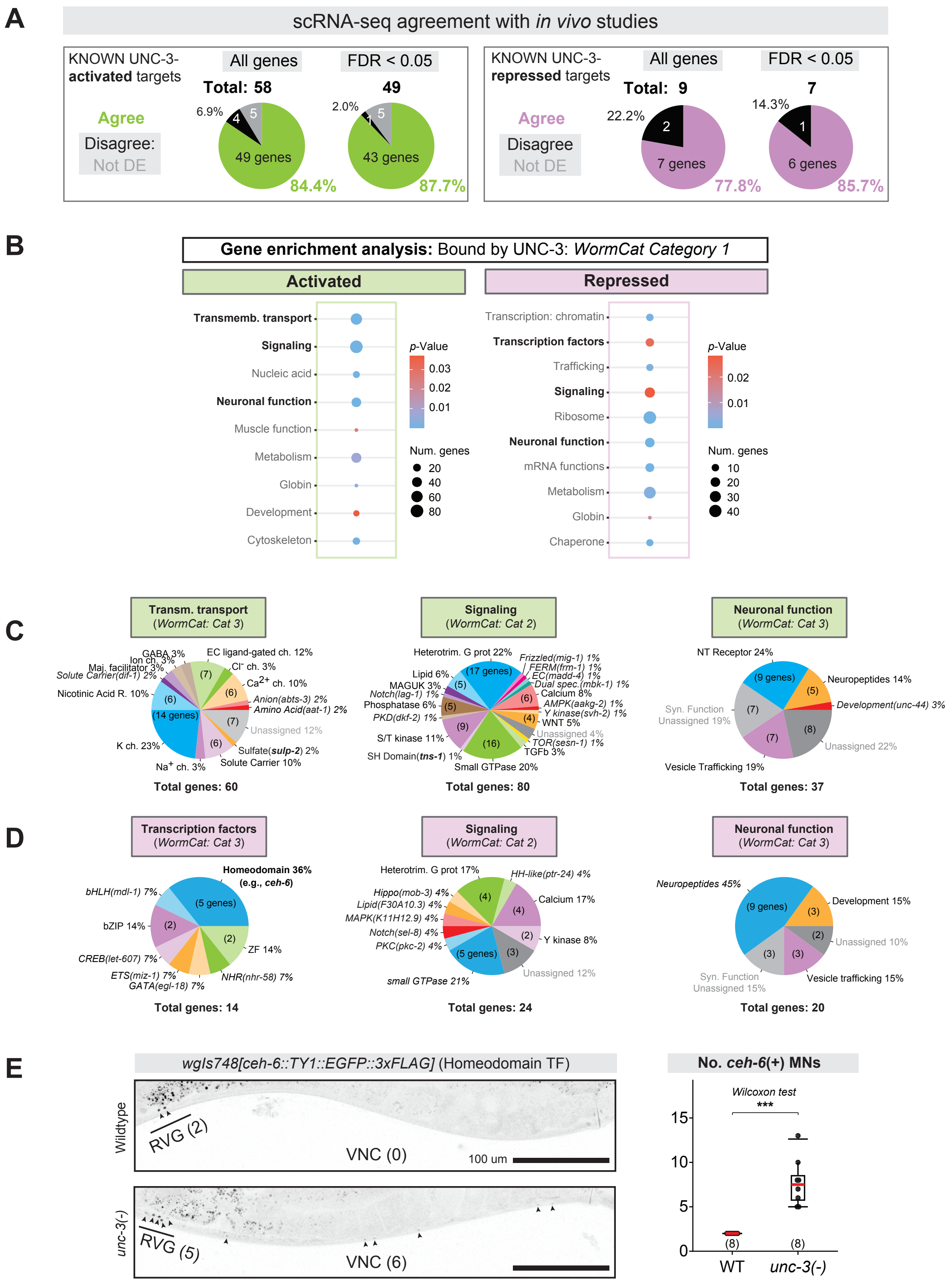


**Figure S2. ScRNA-seq analysis strongly agrees with prior UNC-3/EBF studies. (A)** Pie charts depicting the degree of agreement between scRNA-seq DEGs and UNC-3 targets previously identified in candidate gene studies. For known activated (left) and repressed (right) targets, agreement is provided with and without standard False Discovery Rate (FDR) < 0.05 threshold used throughout this study. **(B)** Dotplots depicting enriched categories (Category resolution 1; WormCat2[60]) in scRNA-seq DEGs. Dot color indicates significant; dot size indicates the number of genes in each category. **(C-D)** Pie charts depicting identity and proportion of activated (C) and repressed (D) DEGs in categories with bold text in C (Category resolution 2; WormCat2[60]). **(E)** Validation of a representative gene (*ceh-6*) in the most enriched TF subcategory, homeodomain, among repressed DEGs. Number of animals (N) is listed on plot. Wilcoxon rank-sum test was selected based on non-normal data; p<.05 = *; p<.01 = **; p<.001 = ***. Box-plot elements: thick horizontal line (red) = median; box = 25th to 75th percentiles (interquartile range); whiskers extend to the furthest data point within 1.5 × IQR from the quartiles (or the min/max if all points lie within this range); individual data points are overlaid as black dots.
