## Supplementary material for "Intrinsic and non-cell autonomous roles for a neurodevelopmental syndrome-linked transcription factor": Figure S3

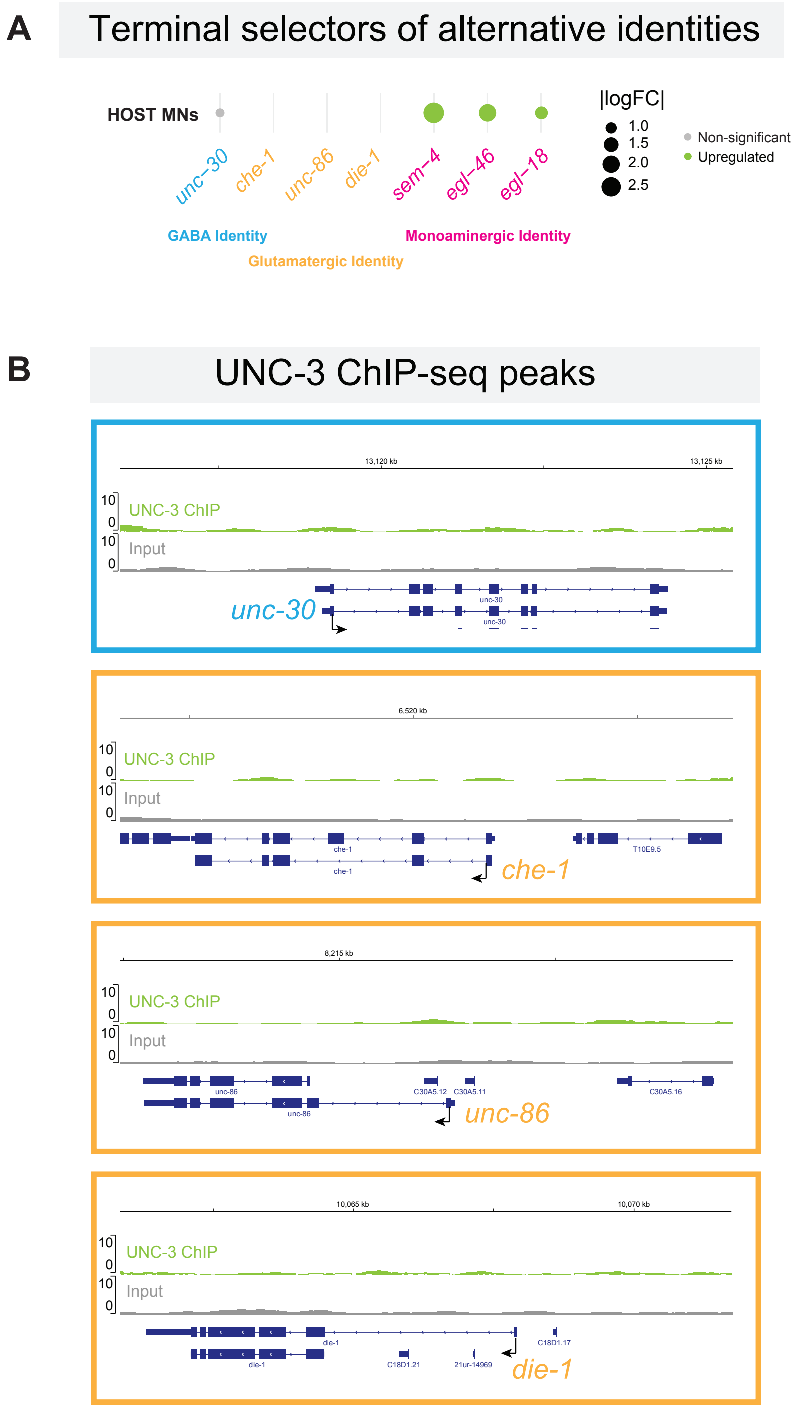


**Figure S3. UNC-3/EBF does not repress selectors of GABA or glutamatergic identity in host MNs. (A)** Dotplot depicting expression of alternative terminal selector TF genes in WT vs. *unc-3(-)* host MNs. We detected no ectopic expression of *unc-30* (GABA) or *che-1*, *unc-86*, and *die-1* (glutamatergic) neuronal identities the mutant. However, selectors of monoaminergic neuronal identities (*sem-4*, *egl-46*, *egl-18*) were upregulated in the absence of *unc-3*, even though monoaminergic NT identity markers (*cat-1, tph-1*) were ectopically expressed in relatively few *unc-3(-)* host MNs. **(B)** UNC-3 ChIP-seq peaks in the cis-regulatory loci of alternative terminal selector genes (*unc-30, che-1, unc-86, die-1*) revealed no significant binding. Tracks were generated using Integrated Genomics Viewer (IGV) (Robinson et al., 2011; Thorvaldsdottir et al., 2012).
