## Supplementary material for "Intrinsic and non-cell autonomous roles for a neurodevelopmental syndrome-linked transcription factor": Figure S4

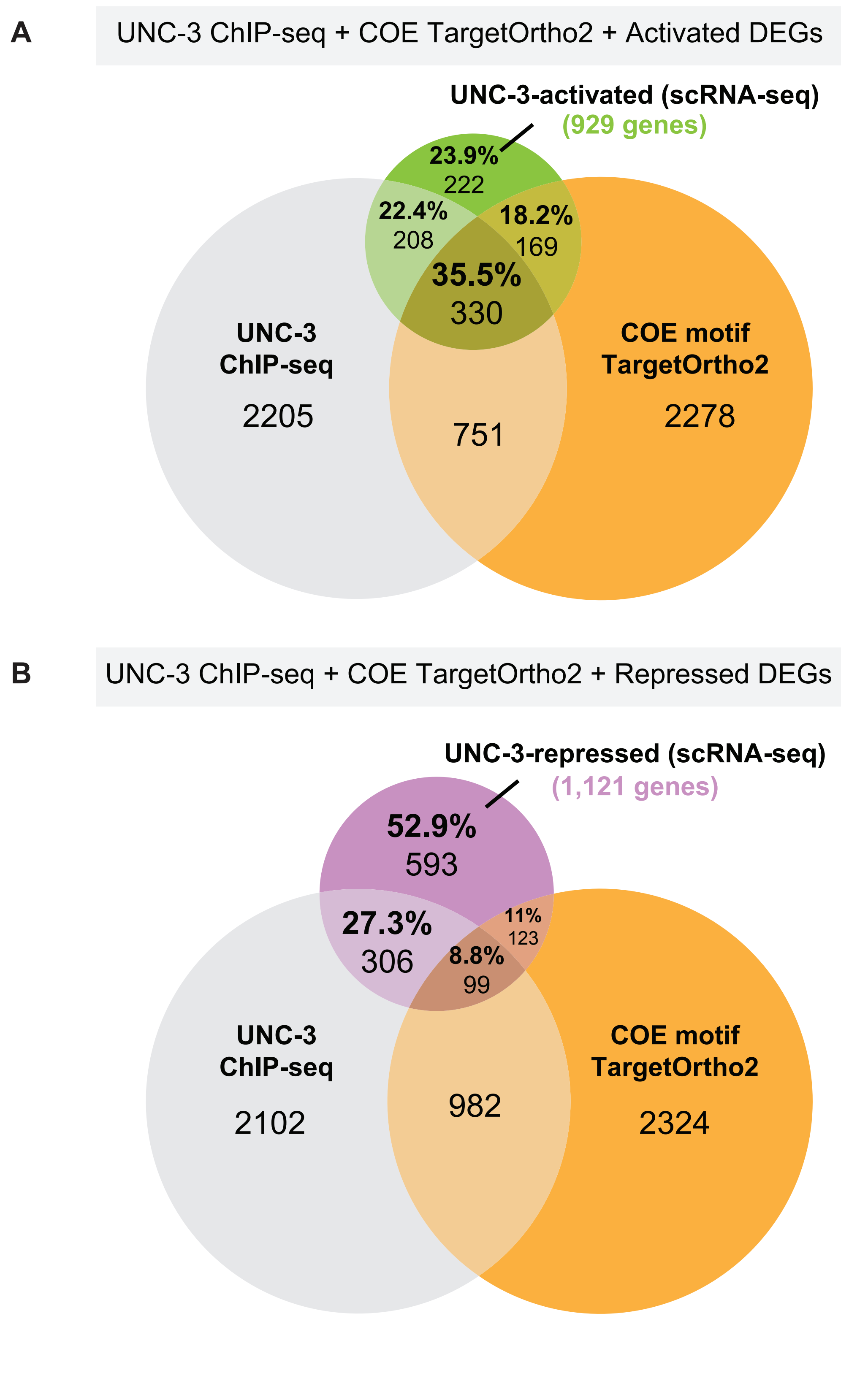


**Figure S4. Integration of genomic and transcriptomic datasets supports COE-independent repression by UNC-3. (A-B)** Venn diagrams depicting the proportion of activated (A) or repressed (B) DEGs shared among UNC-3 ChIP-seq targets (Li et al., 2020), COE TargetOrtho2 predicted targets (Rumley et al., 2025), or both.
