## Supplementary figures and images for "Intrinsic and non-cell autonomous roles for a neurodevelopmental syndrome-linked transcription factor"

### Figure S5

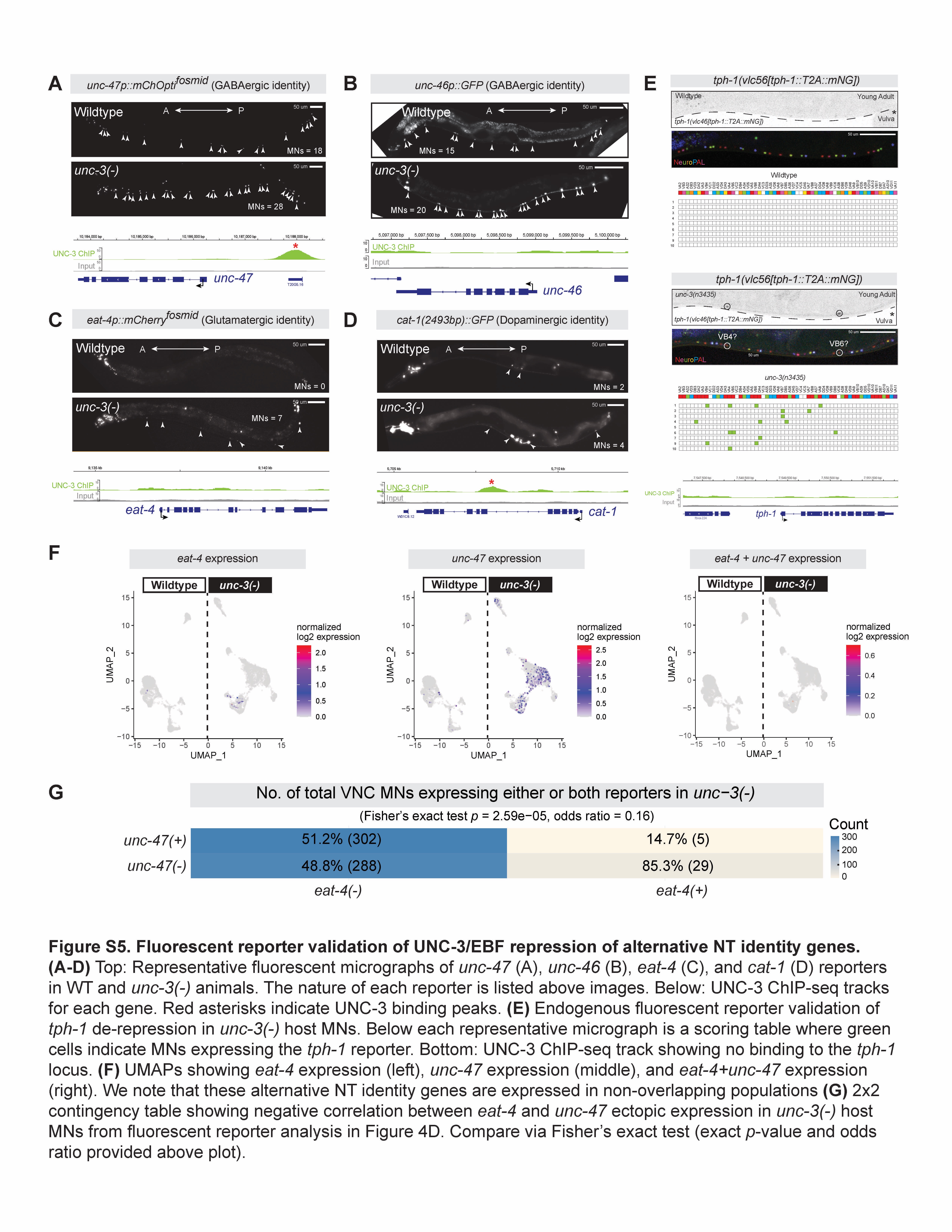
