## Supplementary material for "Intrinsic and non-cell autonomous roles for a neurodevelopmental syndrome-linked transcription factor": Figure S6

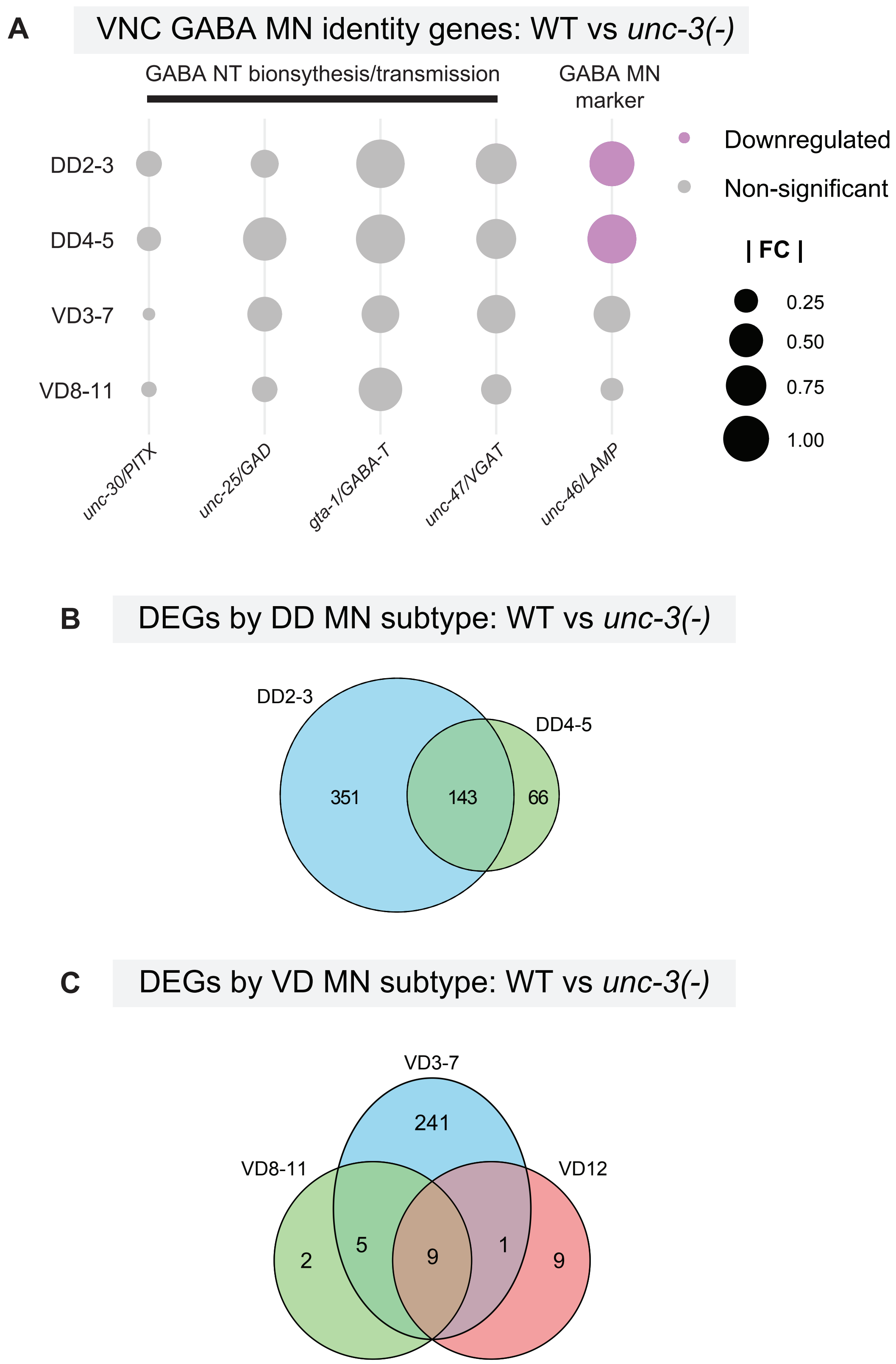


**Figure S6. GABA NT identity is intact in downstream MNs of *unc-3(-)* mutants. (A)** Dot plot depicting no dysregulation of known GABA MN NT identity genes in GABA MN subclasses of *unc-3(-)* animals. However, we note that the GABA MN identity marker *unc-46/LAMP* is downregulated in the absence of *unc-3* (validated in Fig. S8). **(B-C)** Venn diagrams depicting the number of unique and shared DEGs across DD (B) and VD (C) GABA MN subclasses.
