## Supplementary material for "Intrinsic and non-cell autonomous roles for a neurodevelopmental syndrome-linked transcription factor": Figure S7

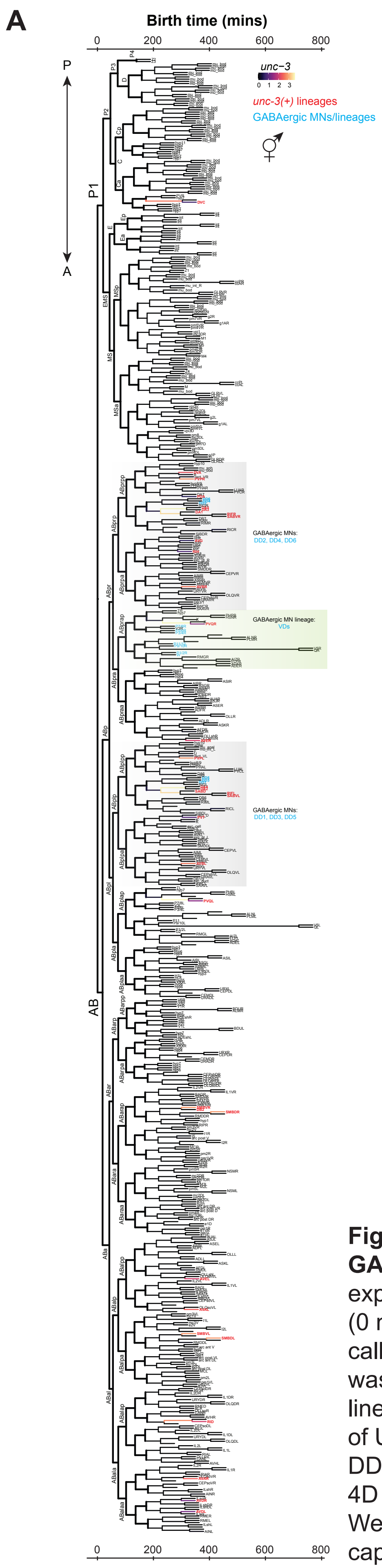


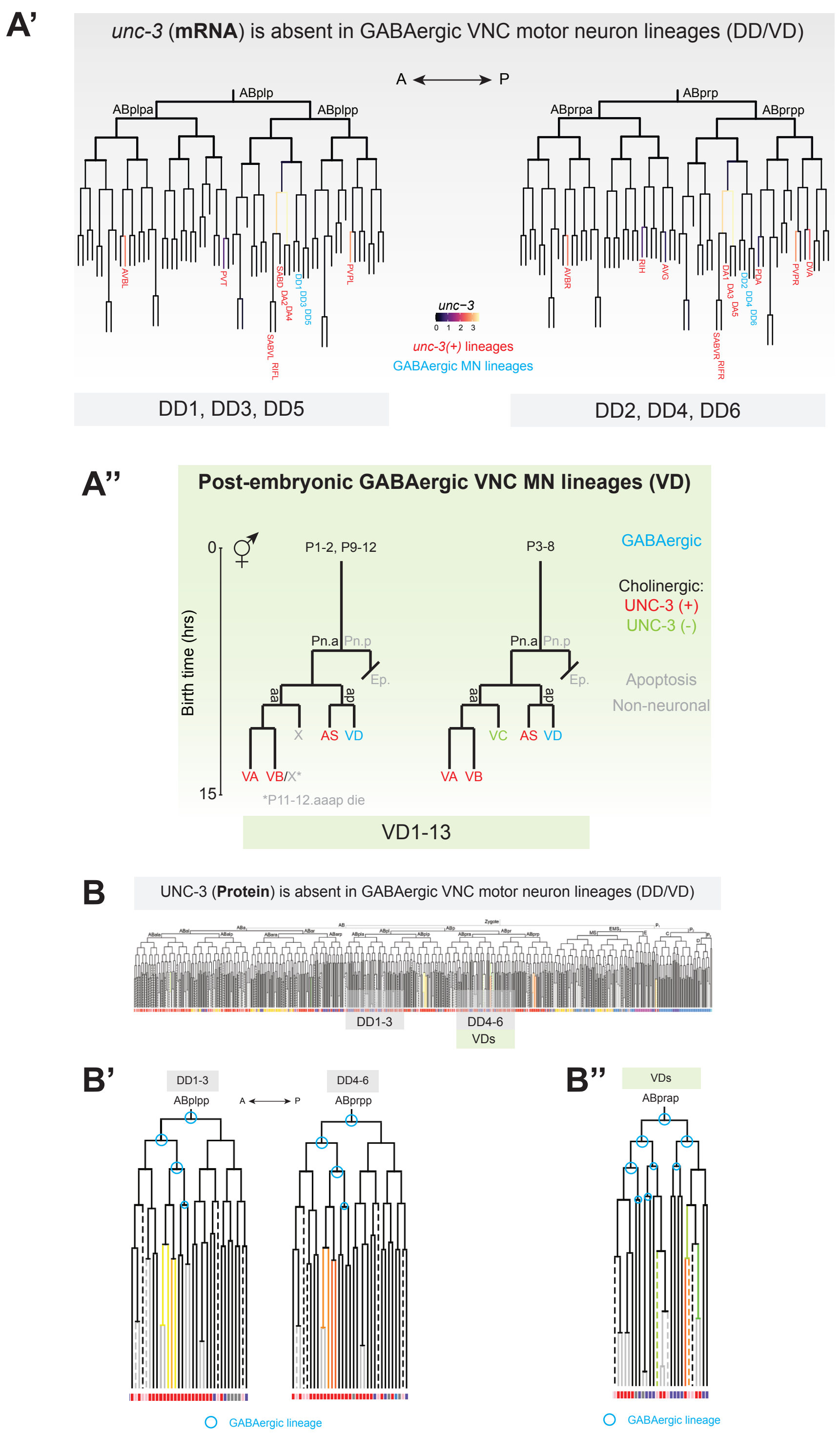


**Figure S7. UNC-3 is not expressed in the lineage of downstream GABA (DD/VD) MNs. (A-A'')** Lineage diagram of *unc-3* (mRNA) expression from scRNA-seq in all embryonic cells (A) from fertilization (0 mins) to the end of embryogenesis (800 minutes post fertilization), with callouts to the lineages that generate the DD (A') and VD (A'') MNs. Tree was generated using VisCell0 (Packer et al., 2019). Cells with *unc-3(+)* lineages are in red, GABA (DD/VD) MNs are in blue. **(B-B'')** Lineage diagram of UNC-3 (protein) expression in *C. elegans* embryogenesis, with callouts to the DD (B') and VD (B'') MN lineages. Tree and UNC-3 expression derive from the 4D single-cell protein atlas of TFs in C. elegans embryogenesis (Ma et al., 2021). We note that while the VD MNs are born post embryonically and thus not captured in these data, their ancestor cells do not express *unc-3* RNA or protein throughout embryogenesis.
