## Supplementary material for "Intrinsic and non-cell autonomous roles for a neurodevelopmental syndrome-linked transcription factor": Figure S8

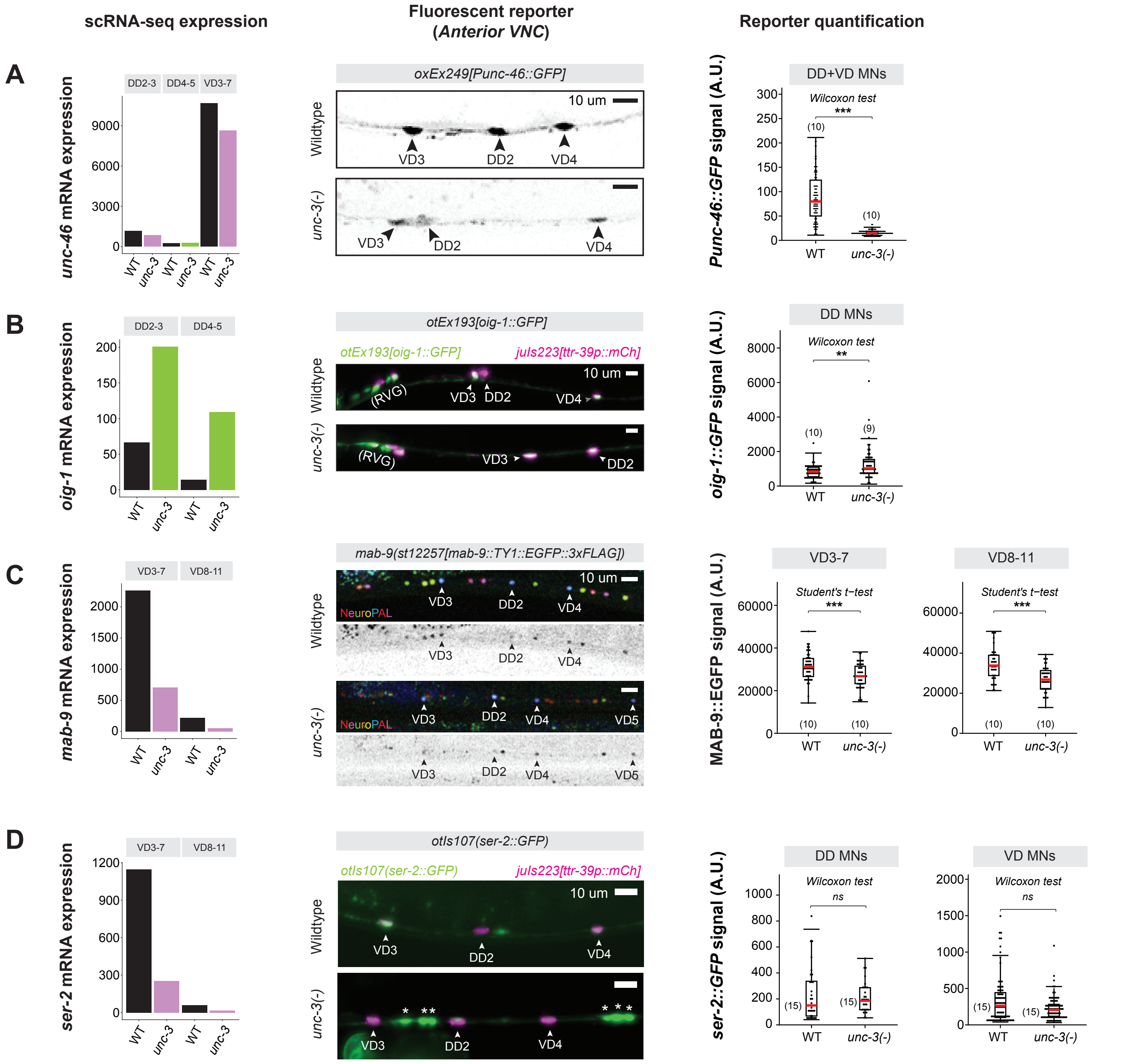


**Figure S8. Fluorescent reporter validation of gene expression changes in downstream MNs of *unc-3(-)* mutants. (A-D)** ScRNA-seq expression (left), representative fluorescent micrographs (middle), and quantifications (right) of UNC-3 non-cell-autonomous effects on expression of *unc-46/LAMP*, (A) *oig-1* (B), *mab-9/TBX20* (C), and *ser-2/HTR1A* (D). **(D)** While scRNA-seq detected downregulation of *ser-2/HTR1A* in downstream MNs of *unc-3(-)* animals, we detected no significant differences between WT and *unc-3(-)* backgrounds using the *otIs107(ser-2::GFP)* fluorescent reporter. Asterisks indicate ectopic expression of *ser-2::GFP* in host MNs. Number of animals (N) is listed in each plot. Statistical tests were selected based on data distribution as follows: Normally distributed data (Shapiro-Wilk p > 0.05 in both groups) with equal variances (Levene's test p > 0.05) = Student's t-test; Normally distributed with unequal variances = Welch's t-test; Non-normal data = Wilcoxon rank-sum test. For all quantifications/statistical tests, p>0.05 = not significant, ns; p<.05 = *; p<.01 = **; p<.001 = ***. Box-plot elements: thick horizontal line (red) = median; box = 25th to 75th percentiles (interquartile range); whiskers extend to the furthest data point within 1.5 × IQR from the quartiles (or the min/max if all points lie within this range); individual data points are overlaid as black dots.
