## Supplementary material for "Intrinsic and non-cell autonomous roles for a neurodevelopmental syndrome-linked transcription factor": Figure S9

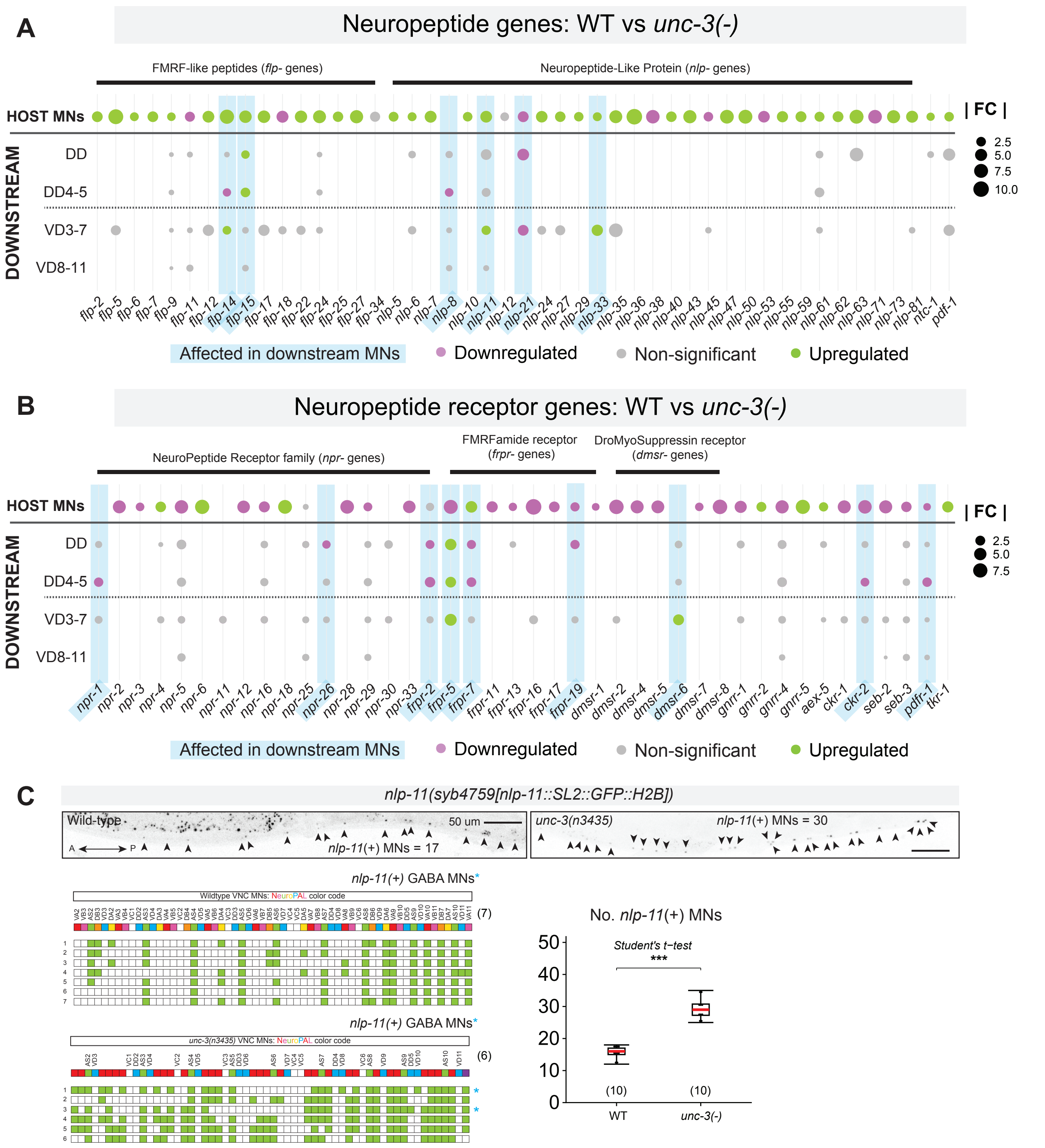


**Figure S9. Neuropeptidergic signaling is broadly dysregulated in host and downstream MNs of *unc-3(-)* animals. (A-B)** Dot plots depicting expression of genes encoding neuropeptides (A) or neuropeptide receptors (B) in WT vs. *unc-3(-)* host and downstream (GABA) MNs. Genes have been grouped into their corresponding families. **(C)** Endogenous fluorescent reporter validation of *nlp-11* dysregulation in host and downstream (GABA) MNs of *unc-3(-)* animals. Top: Fluorescent micrographs of endogenous *nlp-11* expression in WT (left) and *unc-3(-)* (right) backgrounds. Bottom left: single-cell scoring of *nlp-11* fluorescent expression across individual animals harboring the NeuroPAL transgene. Blue asterisks to the right of each row/animal scored indicated ectopic expression in GABA MNs. Bottom right: Dot plot comparing number of *nlp-11(+)* VNC MNs in WT and *unc-3(-)* backgrounds. Quantifications were performed at young adult stage. Bottom left: Number of animals (N) scored at single-cell resolution with NeuroPAL is listed to the left of scoring tables; Bottom right: Number of animals (N) score for *nlp-11(+)* VNC MNs is listed on the plot. Data were compared using a Student's t-test due to normally distributed data (Shapiro-Wilk $p > 0.05$ in both groups) with equal variances (Levene's test $p > 0.05$), $p>.5$ = Not significant, ns; $p<.05$ = *; $p<.01$ = **; $p<.001$ = ***. Box-plot elements: thick horizontal line (red) = median; box = 25th to 75th percentiles (interquartile range); whiskers extend to the furthest data point within 1.5 × IQR from the quartiles (or the min/max if all points lie within this range); individual data points are overlaid as black dots.
