## Supplementary material for "Intrinsic and non-cell autonomous roles for a neurodevelopmental syndrome-linked transcription factor": Figure S10

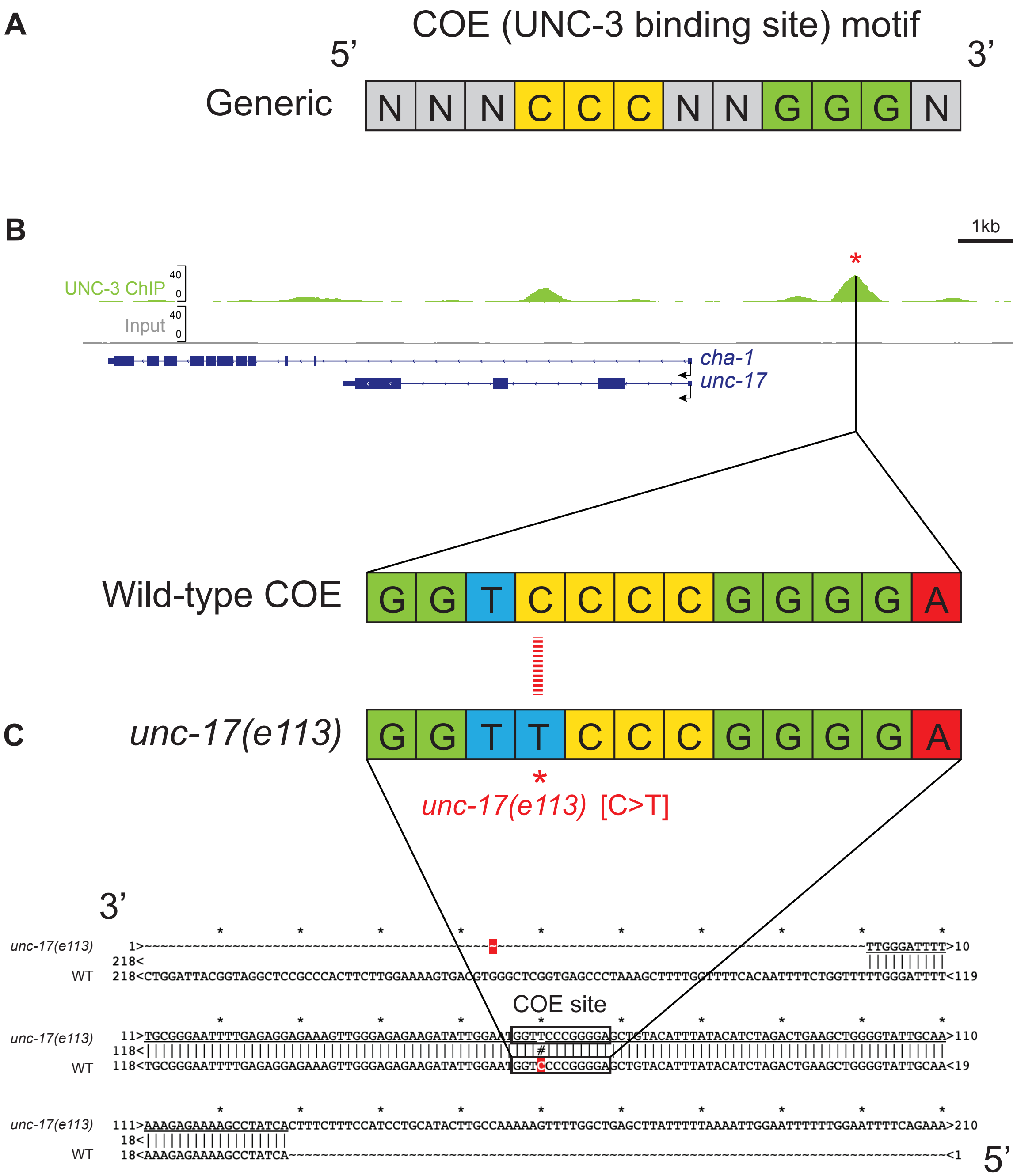


**Figure S10. The *unc-17(e113)* variant targets the COE motif upstream of the *cha-1/unc-17* locus.** **(A)** The sequence of UNC-3’s cognate binding site for activation, the COE motif. The letter N refers to variable identities of nucleotides in that position, whereas all other letters refer to specific nucleotide identities: A = Adenine, T = Thymine, C = Cytosine, G = Guanine. **(B)** Top: UNC-3 ChIP-seq track showing binding in a single COE-containing peak ~2kb upstream of the *cha-1/unc-17* locus. Bottom: exact COE sequenced found with the UNC-3 ChIP-seq peak. (C) Sanger sequencing revealed that the *unc-17(e113)* variant is a C>T point mutation affecting a deeply conserved residue in the COE motif corresponding to the site of UNC-3 binding.
