## Supplementary material for "Intrinsic and non-cell autonomous roles for a neurodevelopmental syndrome-linked transcription factor": Figure S11

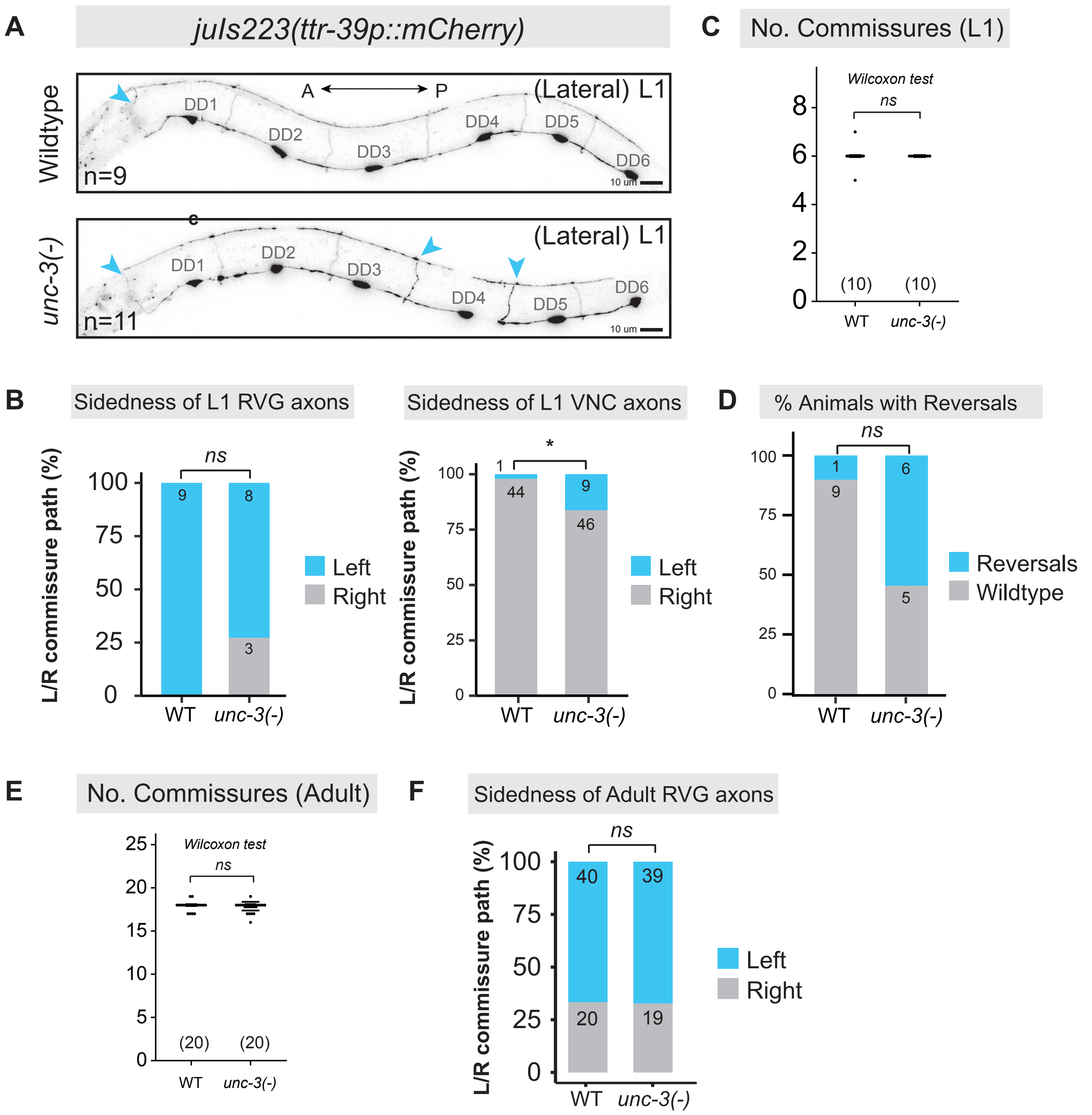


**Figure S11. L1 downstream MNs (DDs) display axon pathfinding defects in the absence of unc-3. (A)** Fluorescence micrographs depicting GABA MN-specific (*ttr-39p::mCherry*) reporter expression in WT (top) and *unc-3(-)* (bottom) backgrounds. Right-to-left commissural reversals are seen from the lateral view in *unc-3(-)* animals. Blue arrowheads = commissures on the left. Number of animals (N) scored is listed on each micrograph. **(B)** Stacked bar chart comparing L1 GABA MN axon reversals in the RVG (left) or VNC (right) of WT and *unc-3(-)* animals. **(C)** Dotplot showing number of GABA MN commissures in L1 animals of each genotype. **(D)** Stacked bar chart showing comparison of the total percentage of L1 animals with at least one GABA MN reversal in each genotype. **(E)** Dotplot showing number of GABA MN commissures in adult animals of each genotype. **(F)** Stacked bar chart showing no difference in the sidedness/reversals of GABA MNs in the adult RVG between genotypes. All statistical comparisons of defect frequencies (stacked bar charts in B, D, F), Compared via Fisher's exact test, ns = not significant; $p<.05$ = *; $p<.01$ = **; $p<.001$ = ***; $p<.00001$ = ****. Commissure counts in C and E were statistically compared via Wilcoxon Rank-Sum Test based on non-normal data, ns = not significant. Number of animals (N) for stacked bar charts is listed in or above each bar and for dot plots is listed below data.
