## Supplementary material for "Intrinsic and non-cell autonomous roles for a neurodevelopmental syndrome-linked transcription factor": Figure S12

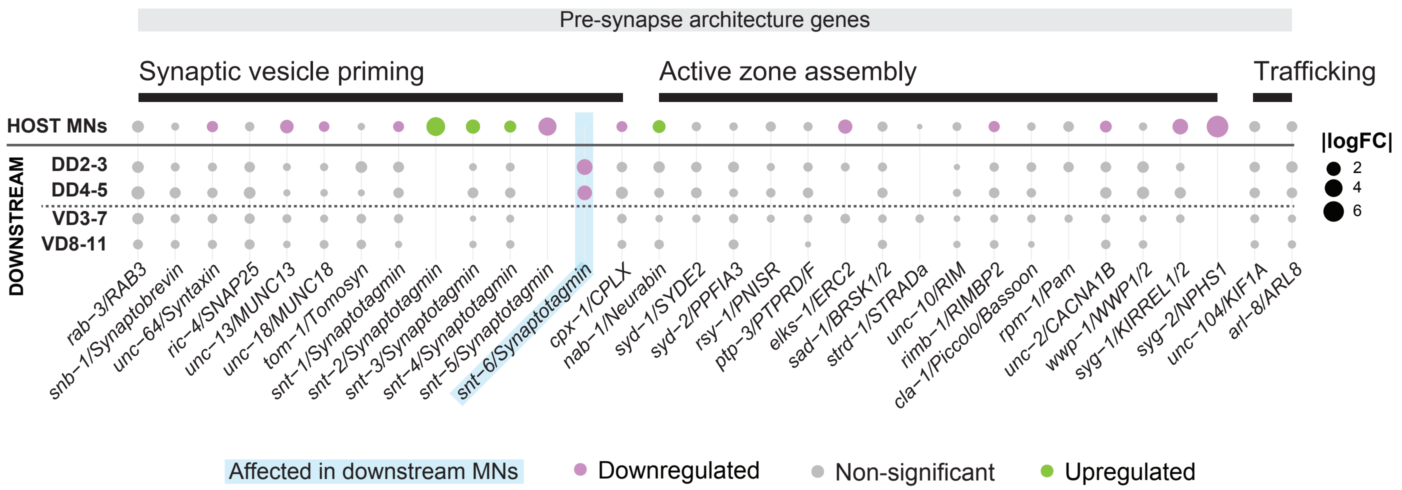


**Figure S12. Expression of presynaptic genes in host and downstream (GABA) MNs in the absence of *unc-3*.** Dot plot depicting DEGs involved in synaptic vesicle priming (left) or active zone assembly (right) in host or downstream MNs.
