## Supplementary material for "Intrinsic and non-cell autonomous roles for a neurodevelopmental syndrome-linked transcription factor": Figure S13

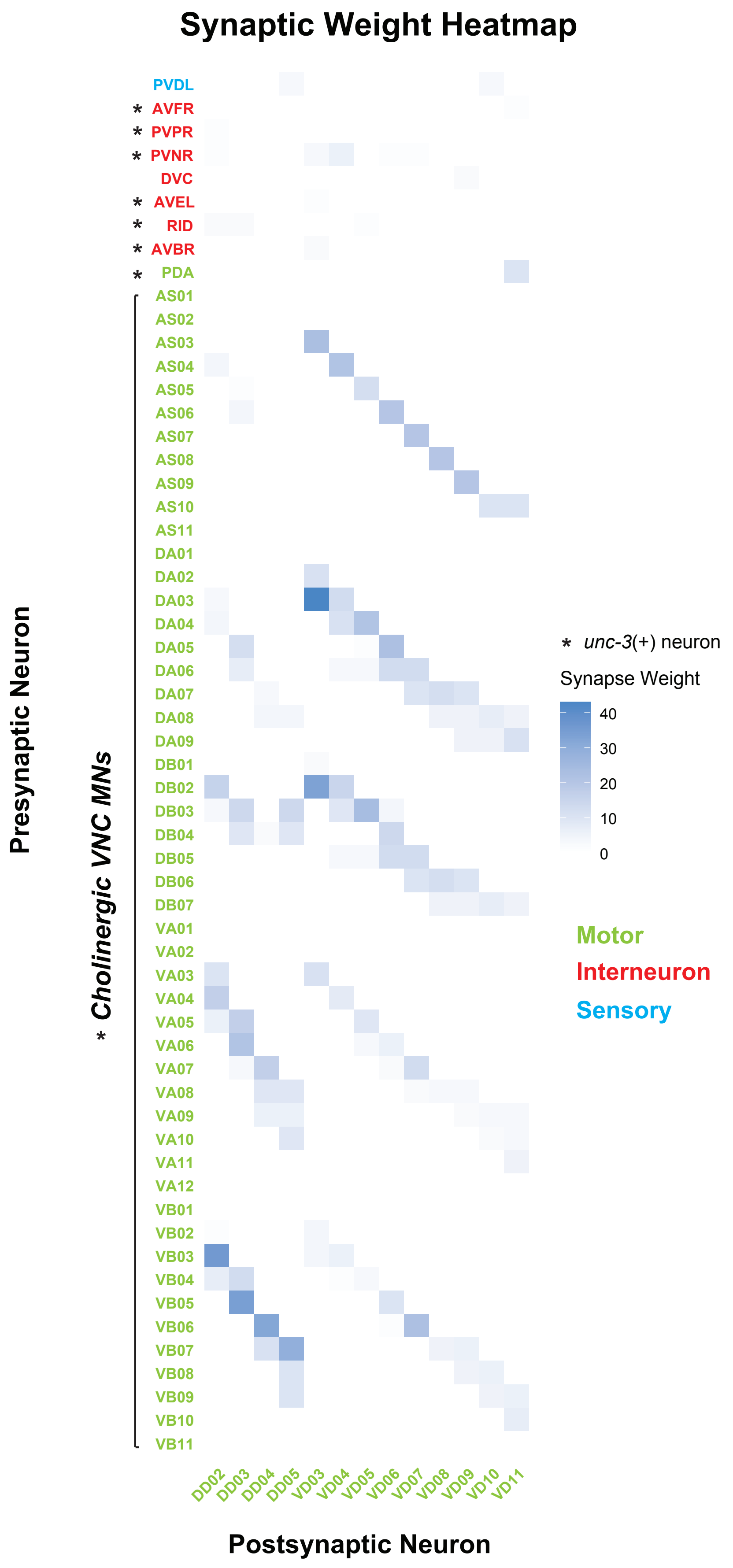


**Figure S13. Downstream (GABA) MNs receive most of their synaptic inputs from host cholinergic MNs.** Heatmap of synaptic weight (i.e., strength) of neurons onto downstream/GABA MNs. Weights are derived from the size of synapses based on electron microscopy reconstructions of the *C. elegans* hermaphrodite nervous system (Cook et al., 2019). We note that the vast majority of synaptic inputs to GABA MNs are from cholinergic MNs, whereas the indicated interneurons (AVFR, PVPR, PVNR, AVEL, RID, AVBR), which are also upstream and *unc-3*-expressing (Taylor et al., 2021), form very few synapses onto GABA MNs of the VNC.
