## Supplementary material for "Intrinsic and non-cell autonomous roles for a neurodevelopmental syndrome-linked transcription factor": Figure S14

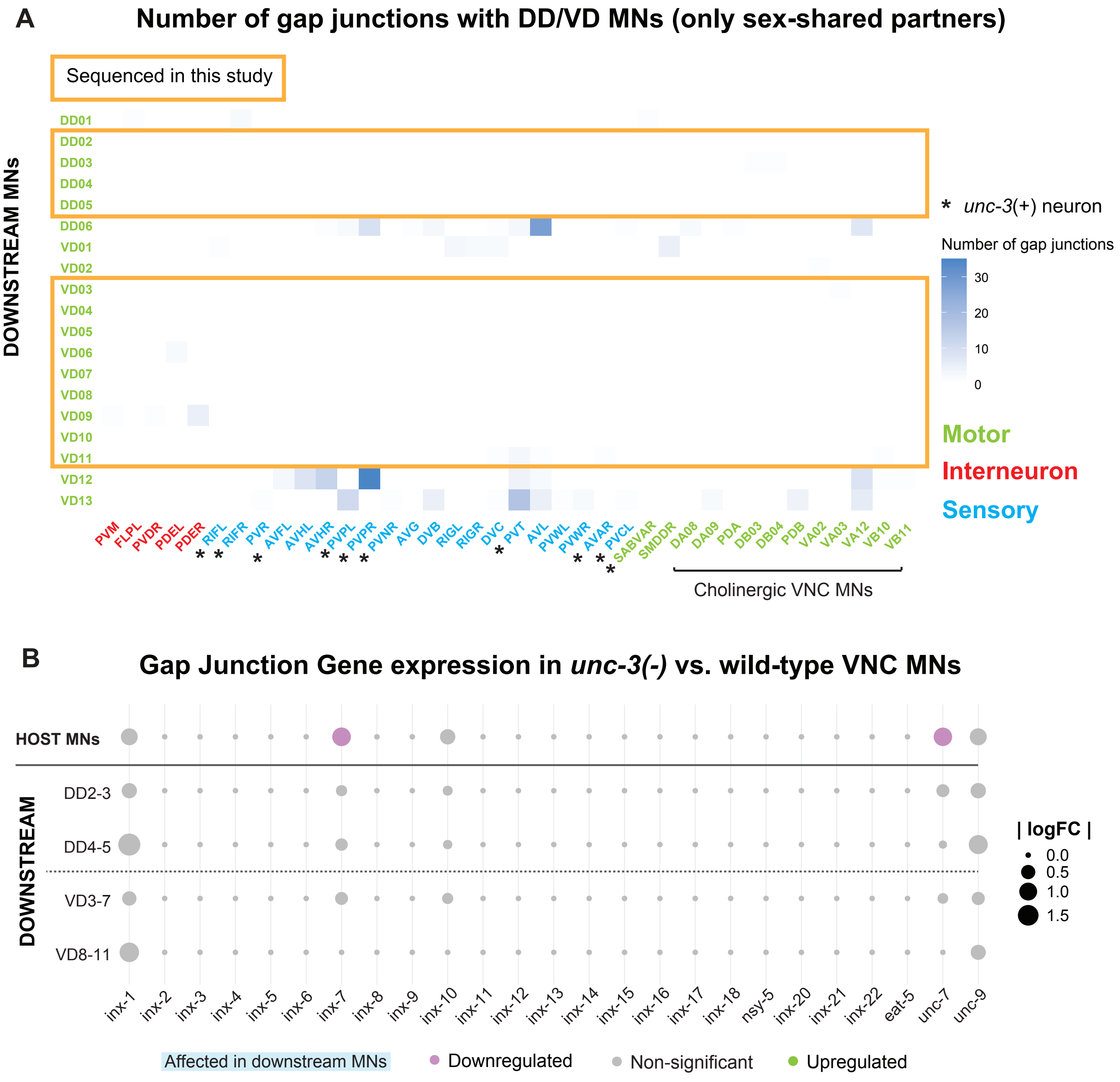


**Figure S14. Downstream (GABA) MNs form almost no gap junctions with nearby cells. (A)** Heatmap depicting the number of gap junctions formed between downstream MNs and other cells based on electron microscopy reconstructions of the *C. elegans* hermaphrodite nervous system (Cook et al., 2019). GABA MNs sequenced in this study are contained within orange boxes. We note that GABA MNs of the VNC form almost no gap junctions with host MNs. Neuron labels are colored by functional category: MNs = green, Interneurons = Red, sensory neurons = Blue. Asterisks indicate *unc-3(+)* neuron classes. **(B)** Dot plot depicting differential expression of gap junction genes (i.e., innexins) in host or downstream MNs. We note that no gap junction genes were impacted in GABA MNs of *unc-3(-)* MNs. Dot size indicates absolute value of the log fold change (|logFC|) in WT vs. *unc-3(-)*.
