## Supplementary material for "Intrinsic and non-cell autonomous roles for a neurodevelopmental syndrome-linked transcription factor": Figure S15

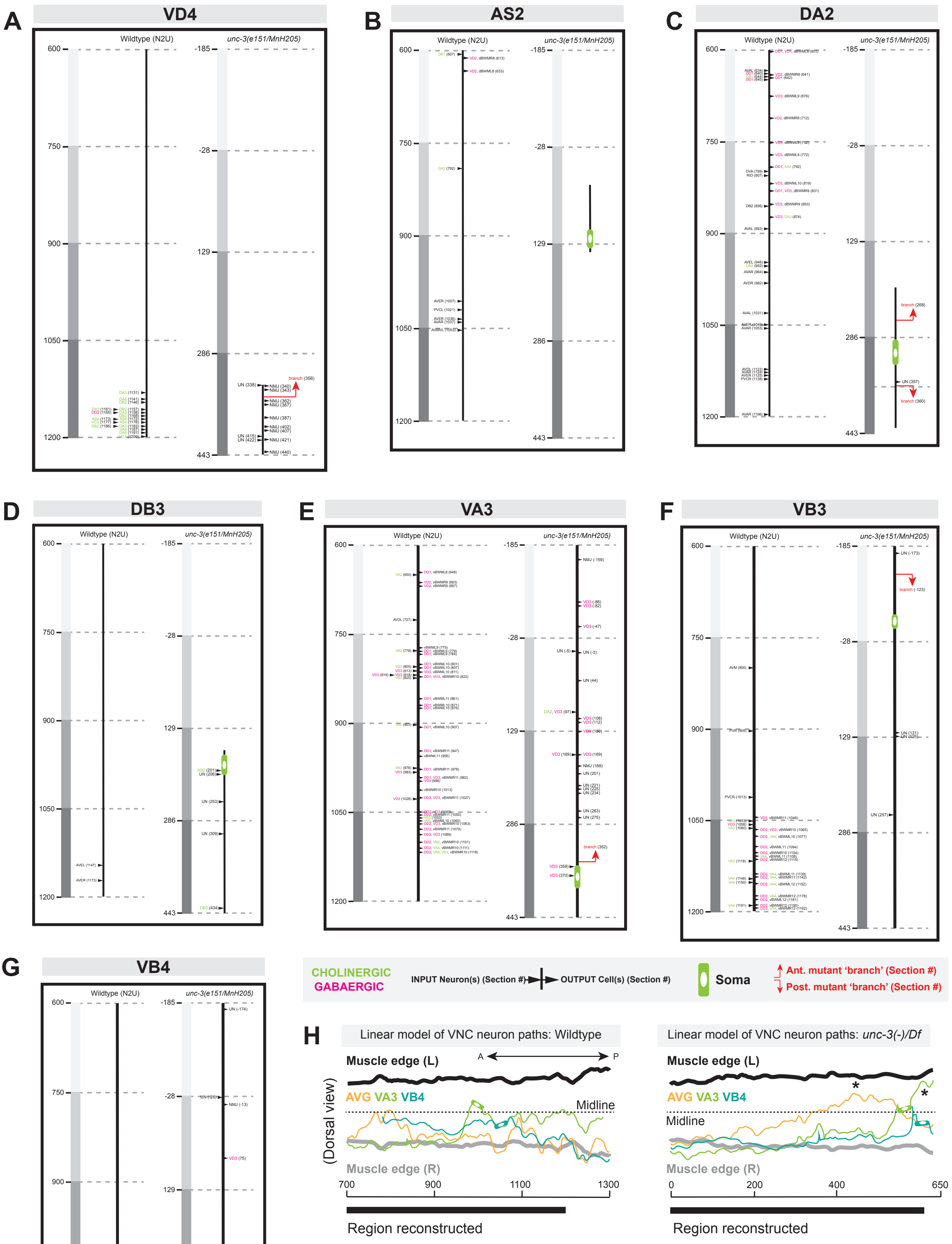


**Figure S15. Electron microscopy reconstructions of host and down-stream MNs. (A-G)** Synaptic inputs (left of axon/soma) and outputs (right of axon/soma) of the VD4 GABA (A) and AS2 (B), DA2 (C), DB3 (D), VA3 (E), VB3 (F), VB4 (G) host MNs in the WT and *unc-3(-)/Df* backgrounds. **(H)** Linear tracing of individual neurons, including the AVG interneuron and VA3 and VB4 host cholinergic MNs, revealed inappropriate midline crossings (asterisks), an aberrant behavior not observed in its WT counterpart.
