## Supplementary material for "Intrinsic and non-cell autonomous roles for a neurodevelopmental syndrome-linked transcription factor": File S5

### Homer *de novo* Motif Results

#### Downregulated Genes

Total target sequences = 650
Total background sequences = 41701
* - possible false positive

| Rank | Motif | P-value | log P-pvalue | % of Targets | % of Background | STD(Bg STD) | Best Match |
|---|---|---|---|---|---|---|---|
| 1 | 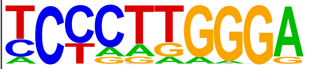 | 1e-40 | -9.390e+01 | 8.92% | 0.77% | 18.2bp (23.5bp) | POL013.1_MED-1/Jaspar(0.502) |
| 2 | 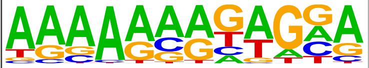 | 1e-21 | -4.950e+01 | 30.92% | 15.73% | 20.7bp (28.0bp) | blmp-1/MA0537.1/Jaspar(0.639) |
| 3 | 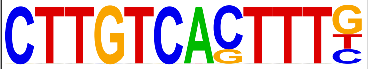 | 1e-13 | -3.155e+01 | 0.92% | 0.00% | 19.9bp (9.8bp) | blmp-1/MA0537.1/Jaspar(0.727) |
| 4 * | 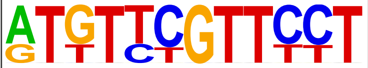 | 1e-11 | -2.560e+01 | 0.77% | 0.00% | 11.5bp (11.2bp) | MF0009.1_TRP(MYB)_class/Jaspar(0.638) |
| 5 * | 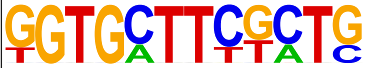 | 1e-11 | -2.560e+01 | 0.77% | 0.00% | 15.8bp (7.7bp) | POL008.1_DCE_S_I/Jaspar(0.564) |
| 6 * | 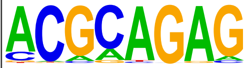 | 1e-10 | -2.309e+01 | 14.00% | 6.83% | 19.7bp (28.4bp) | eor-1/MA0543.1/Jaspar(0.659) |
| 7 * | 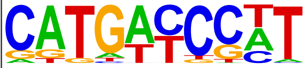 | 1e-9 | -2.163e+01 | 2.77% | 0.41% | 20.5bp (22.3bp) | MF0004.1_Nuclear_Receptor_class/Jaspar(0.662) |
| 8 * | 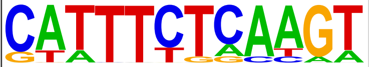 | 1e-9 | -2.123e+01 | 2.46% | 0.32% | 17.7bp (23.1bp) | ceh-22/MA0264.1/Jaspar(0.627) |
| 9 * | 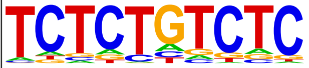 | 1e-9 | -2.097e+01 | 16.77% | 9.18% | 18.0bp (37.6bp) | eor-1/MA0543.1/Jaspar(0.803) |
| 10 * | 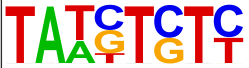 | 1e-8 | -2.070e+01 | 20.92% | 12.47% | 18.9bp (26.0bp) | unc-62/MA0918.1/Jaspar(0.576) |
| 11 * | 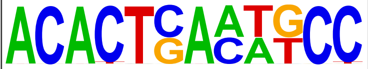 | 1e-8 | -2.013e+01 | 0.77% | 0.01% | 11.8bp (12.6bp) | MF0011.1_HMG_class/Jaspar(0.509) |
| 12 * | 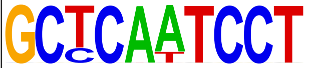 | 1e-8 | -1.982e+01 | 0.62% | 0.00% | 19.7bp (4.2bp) | unc-30/MA1443.1/Jaspar(0.610) |
| 13 * | 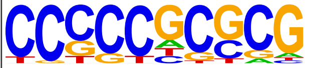 | 1e-8 | -1.916e+01 | 4.77% | 1.38% | 16.1bp (27.4bp) | dpy-27/MA0540.1/Jaspar(0.578) |
| 14 * | 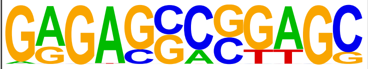 | 1e-8 | -1.876e+01 | 2.62% | 0.43% | 16.9bp (25.3bp) | POL013.1_MED-1/Jaspar(0.594) |
| 15 * | 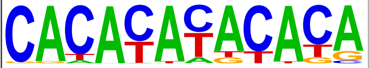 | 1e-7 | -1.814e+01 | 3.69% | 0.91% | 18.2bp (27.6bp) | daf-12/MA0538.1/Jaspar(0.755) |
| 16 * | 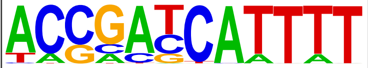 | 1e-7 | -1.813e+01 | 1.54% | 0.12% | 16.9bp (26.0bp) | POL002.1_INR/Jaspar(0.541) |
| 17 * | 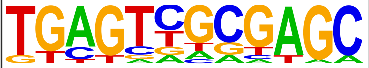 | 1e-7 | -1.786e+01 | 2.46% | 0.40% | 21.1bp (20.8bp) | POL010.1_DCE_S_III/Jaspar(0.602) |
| 18 * | 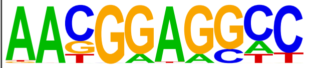 | 1e-7 | -1.760e+01 | 0.77% | 0.01% | 23.3bp (15.8bp) | POL013.1_MED-1/Jaspar(0.635) |

| | | | | | | | |
|---|---|---|---|---|---|---|---|
| | | | | | | (32.9bp) | |
| 20 * | 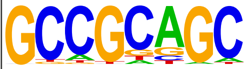 | 1e-6 | -1.468e+01 | 5.08% | 1.87% | 16.7bp (23.6bp) | POL013.1_MED-1/Jaspar(0.640) |
| 21 * | 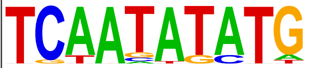 | 1e-6 | -1.466e+01 | 1.38% | 0.14% | 18.9bp (21.1bp) | MF0008.1_MADS_class/Jaspar(0.670) |
| 22 * | 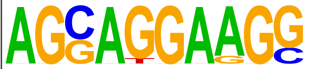 | 1e-5 | -1.358e+01 | 1.23% | 0.12% | 16.3bp (21.3bp) | POL008.1_DCE_S_I/Jaspar(0.661) |
| 23 * | 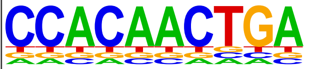 | 1e-5 | -1.221e+01 | 0.46% | 0.01% | 20.5bp (14.2bp) | hlh-1/MA0545.1/Jaspar(0.632) |
| 24 * | 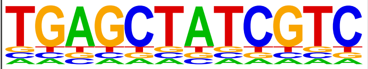 | 1e-5 | -1.221e+01 | 0.46% | 0.01% | 5.4bp (12.2bp) | nhr-6/MA1451.1/Jaspar(0.518) |
| 25 * | 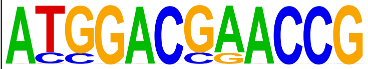 | 1e-5 | -1.221e+01 | 0.46% | 0.01% | 14.9bp (5.3bp) | MF0009.1_TRP(MYB)_class/Jaspar(0.557) |
| 26 * | 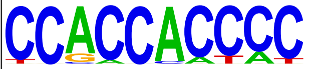 | 1e-5 | -1.195e+01 | 1.69% | 0.30% | 15.0bp (28.1bp) | POL003.1_GC-box/Jaspar(0.617) |
| 27 * | 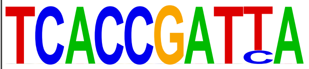 | 1e-4 | -1.015e+01 | 0.46% | 0.01% | 17.9bp (13.6bp) | ceh-28/MA1445.1/Jaspar(0.606) |
| 28 * | 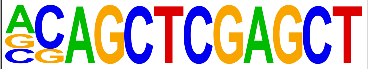 | 1e-3 | -9.017e+00 | 0.31% | 0.00% | 16.7bp (0.0bp) | POL010.1_DCE_S_III/Jaspar(0.544) |
| 29 * | 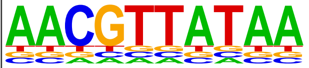 | 1e-3 | -7.641e+00 | 0.31% | 0.01% | 22.0bp (10.8bp) | zip-8/MA1704.1/Jaspar(0.622) |
| 30 * | 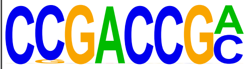 | 1e-2 | -6.118e+00 | 0.92% | 0.20% | 13.9bp (23.9bp) | POL011.1_XCPE1/Jaspar(0.535) |

### Homer Known Motif Enrichment Results

#### Downregulated Genes

Total Target Sequences = 650, Total Background Sequences = 41501

| Rank | Motif | Name | P-value | log P-pvalue | q-value (Benjamini) | # Target Sequences with Motif | % of Targets Sequences with Motif | # Background Sequences with Motif | % of Background Sequences with Motif |
|---|---|---|---|---|---|---|---|---|---|
| 1 | 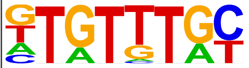 | PHA-4(Forkhead)/cElegans-Embryos-PHA4-ChIP-Seq(modEncode)/Homer | 1e-3 | -7.413e+00 | 0.0066 | 105.0 | 16.15% | 4897.1 | 11.80% |
| 2 | 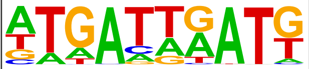 | LIN-39(Homeobox)/cElegans.L3-LIN39-ChIP-Seq(modEncode)/Homer | 1e-2 | -4.978e+00 | 0.0379 | 58.0 | 8.92% | 2646.8 | 6.38% |
