## Supplementary material for "Intrinsic and non-cell autonomous roles for a neurodevelopmental syndrome-linked transcription factor": File S6

### Homer *de novo* Motif Results

#### Upregulated Genes

Total target sequences = 412
Total background sequences = 38500
* - possible false positive

| Rank | Motif | P-value | log P-pvalue | % of Targets | % of Background | STD(Bg STD) | Best Match/Details |
|---|---|---|---|---|---|---|---|
| 1 | 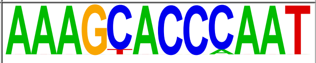 | 1e-20 | -4.761e+01 | 2.67% | 0.02% | 17.6bp (13.8bp) | ceh-10::ttx-3/MA0263.1/Jaspar(0.622) |
| 2 | 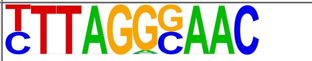 | 1e-19 | -4.376e+01 | 2.91% | 0.04% | 16.9bp (23.8bp) | MF0006.1_bZIP_cEBP-like_subclass/Jaspar(0.668) |
| 3 | 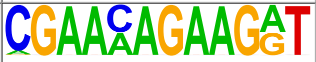 | 1e-18 | -4.371e+01 | 2.67% | 0.02% | 17.6bp (14.2bp) | POL008.1_DCE_S_I/Jaspar(0.583) |
| 4 | 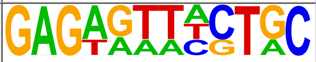 | 1e-18 | -4.267e+01 | 2.67% | 0.03% | 21.0bp (25.1bp) | daf-12/MA0538.1/Jaspar(0.558) |
| 5 | 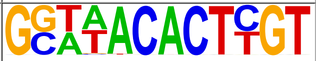 | 1e-16 | -3.694e+01 | 1.94% | 0.01% | 16.3bp (17.7bp) | daf-16/MA1446.1/Jaspar(0.580) |
| 6 | 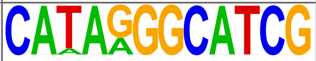 | 1e-15 | -3.640e+01 | 1.70% | 0.01% | 13.5bp (21.5bp) | PL0012.1_hlh-2::hlh-8/Jaspar(0.534) |
| 7 | 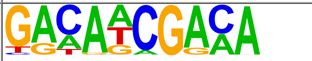 | 1e-14 | -3.324e+01 | 5.83% | 0.68% | 18.3bp (24.6bp) | blmp-1/MA0537.1/Jaspar(0.669) |
| 8 | 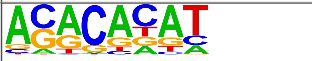 | 1e-14 | -3.305e+01 | 33.01% | 17.15% | 19.7bp (27.2bp) | sma-4/MA0925.1/Jaspar(0.662) |
| 9 | 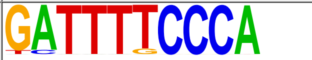 | 1e-13 | -3.058e+01 | 2.91% | 0.11% | 22.4bp (21.3bp) | MF0003.1_REL_class/Jaspar(0.642) |
| 10 | 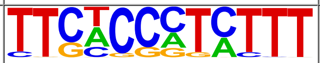 | 1e-12 | -2.978e+01 | 4.13% | 0.33% | 21.2bp (21.5bp) | eor-1/MA0543.1/Jaspar(0.579) |
| 11 | 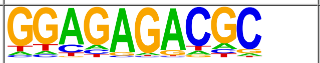 | 1e-12 | -2.970e+01 | 12.14% | 3.60% | 20.2bp (29.4bp) | eor-1/MA0543.1/Jaspar(0.657) |
| 12 * | 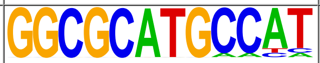 | 1e-11 | -2.632e+01 | 1.46% | 0.01% | 10.4bp (17.2bp) | PL0018.1_hlh-25/Jaspar(0.593) |
| 13 * | 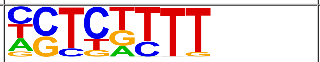 | 1e-9 | -2.287e+01 | 31.55% | 18.45% | 20.0bp (29.6bp) | blmp-1/MA0537.1/Jaspar(0.636) |
| 14 * | 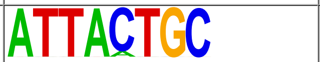 | 1e-9 | -2.117e+01 | 2.91% | 0.24% | 21.6bp (23.7bp) | MF0010.1_Homeobox_class/Jaspar(0.685) |
| 15 * | 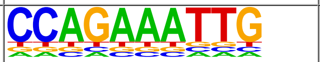 | 1e-8 | -2.031e+01 | 1.46% | 0.03% | 20.5bp (14.6bp) | sma-4/MA0925.1/Jaspar(0.692) |
| 16 * | 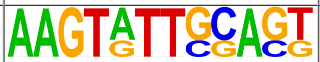 | 1e-8 | -1.846e+01 | 1.21% | 0.02% | 15.1bp (11.5bp) | MF0010.1_Homeobox_class/Jaspar(0.596) |
| 17 * | 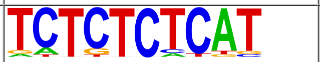 | 1e-7 | -1.719e+01 | 6.07% | 1.63% | 24.8bp (34.9bp) | blmp-1/MA0537.1/Jaspar(0.581) |
| 18 * | 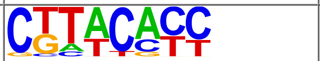 | 1e-7 | -1.690e+01 | 8.98% | 3.26% | 19.5bp (25.2bp) | SD0001.1_at_AC_acceptor/Jaspar(0.671) |
| 19 * | 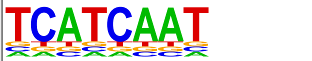 | 1e-7 | -1.685e+01 | 2.91% | 0.36% | 20.4bp (19.5bp) | LIN-39(Homeobox)/cElegans.L3-LIN39-ChIP-Seq(modEncode)/Homer(0.726) |
| 20 * | 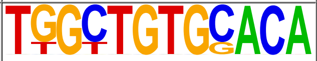 | 1e-6 | -1.580e+01 | 0.73% | 0.00% | 8.7bp (0.0bp) | POL009.1_DCE_S_II/Jaspar(0.603) |
| 21 * | 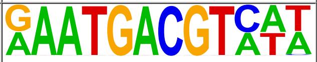 | 1e-6 | -1.410e+01 | 1.21% | 0.04% | 20.2bp (20.1bp) | atf-7/MA1438.1/Jaspar(0.865) |
| 22 * | 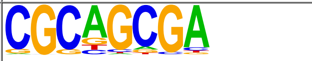 | 1e-5 | -1.322e+01 | 7.04% | 2.60% | 17.9bp (28.3bp) | dpy-27/MA0540.1/Jaspar(0.656) |
| 23 * | 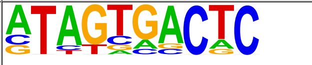 | 1e-5 | -1.318e+01 | 2.67% | 0.42% | 17.9bp (22.8bp) | fos-1/MA1448.1/Jaspar(0.684) |
| 24 * | 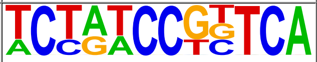 | 1e-5 | -1.234e+01 | 0.97% | 0.03% | 18.1bp (12.5bp) | POL004.1_CCAAT-box/Jaspar(0.571) |
| 25 * | 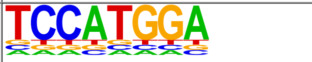 | 1e-4 | -1.136e+01 | 1.21% | 0.07% | 20.2bp (13.7bp) | ceh-48/MA0921.1/Jaspar(0.495) |

### Homer Known Motif Enrichment Results

#### Upregulated Genes

Total Target Sequences = 412, Total Background Sequences = 43733

| Rank | Motif | Name | P-value | log P-pvalue | q-value (Benjamini) | # Target Sequences with Motif | % of Targets Sequences with Motif | # Background Sequences with Motif | % of Background Sequences with Motif |
|---|---|---|---|---|---|---|---|---|---|
| 1 | 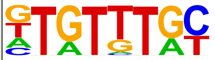 | PHA-4(Forkhead)/cElegans-Embryos-PHA4-ChIP-Seq(modEncode)/Homer | 1e-3 | -8.003e+00 | 0.0037 | 79.0 | 19.17% | 5733.4 | 13.11% |
| 2 | 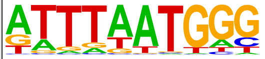 | EGL-5(Homeobox)/cElegans-L3-EGL5-ChIP-Seq(modEncode)/Homer | 1e-3 | -7.941e+00 | 0.0037 | 50.0 | 12.14% | 3208.9 | 7.33% |
| 3 | 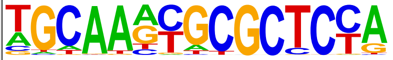 | EFL-1(E2F)/cElegans-L1-EFL1-ChIP-Seq(modEncode)/Homer | 1e-3 | -7.816e+00 | 0.0037 | 11.0 | 2.67% | 334.6 | 0.76% |
| 4 | 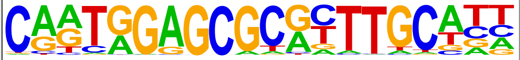 | LIN-15B(Zf)/cElegans-L3-LIN15B-ChIP-Seq(modEncode)/Homer | 1e-3 | -7.811e+00 | 0.0037 | 10.0 | 2.43% | 281.3 | 0.64% |
